# RTX-KG2: a system for building a semantically standardized knowledge graph for translational biomedicine

**DOI:** 10.1101/2021.10.17.464747

**Authors:** E. C. Wood, Amy K. Glen, Lindsey G. Kvarfordt, Finn Womack, Liliana Acevedo, Timothy S. Yoon, Chunyu Ma, Veronica Flores, Meghamala Sinha, Yodsawalai Chodpathumwan, Arash Termehchy, Jared C. Roach, Luis Mendoza, Andrew S. Hoffman, Eric W. Deutsch, David Koslicki, Stephen A. Ramsey

## Abstract

**Background:** Biomedical translational science is increasingly using computational reasoning on repositories of structured knowledge (such as UMLS, SemMedDB, ChEMBL, Reactome, DrugBank, and SMPDB in order to facilitate discovery of new therapeutic targets and modalities. The NCATS Biomedical Data Translator project is working to federate autonomous reasoning agents and knowledge providers within a distributed system for answering translational questions. Within that project and the broader field, there is a need for a framework that can efficiently and reproducibly build an integrated, standards-compliant, and comprehensive biomedical knowledge graph that can be downloaded in standard serialized form or queried via a public application programming interface (API).

**Results:** To create a *knowledge provider* system within the Translator project, we have developed RTX-KG2, an open-source software system for building—and hosting a web API for querying—a biomedical knowledge graph that uses an Extract-Transform-Load (ETL) approach to integrate 70 knowledge sources (including the aforementioned core six sources) into a knowledge graph with provenance information including (where available) citations. The semantic layer and schema for RTX-KG2 follow the standard Biolink model to maximize interoperability. RTX-KG2 is currently being used by multiple Translator reasoning agents, both in its downloadable form and via its SmartAPI-registered interface. Serializations of RTX-KG2 are available for download in both the pre-canonicalized form and in canonicalized form (in which synonyms are merged). The current canonicalized version (KG2.7.3) of RTX-KG2 contains 6.4M nodes and 39.3M edges with a hierarchy of 77 relationship types from Biolink.

**Conclusion:** RTX-KG2 is the first knowledge graph that integrates UMLS, SemMedDB, ChEMBL, DrugBank, Reactome, SMPDB, and 64 additional knowledge sources within a knowledge graph that conforms to the Biolink standard for its semantic layer and schema. RTX-KG2 is publicly available for querying via its API at arax.rtx.ai/api/rtxkg2/v1.2/openapi.json. The code to build RTX-KG2 is publicly available at [github:RTXteam/RTX-KG2](https://github.com/RTXteam/RTX-KG2).

## 1 Background

In translational biomedicine, there is a longstanding need to integrate structured knowledge as a substrate for computational reasoning [1], such as for drug repositioning or finding new therapies for monogenic disorders. Efforts to define a *lingua franca* for a computable and comprehensive biomedical knowledge graph have seen a pivot from controlled vocabularies [2–6] (and their integration in the Unified Medical Language System (UMLS) Metathesaurus [7]) to ontologies in standardized computable representations [8–10]. The World Wide Web has fueled the development of online knowledge-bases updated by literature curation teams, such as KEGG [11], PubChem [12], DrugBank [13], ChEMBL [14], the UniProt Knowledgebase (UniProtKB) [15], the Small Molecule Pathway Database (SMPDB) [16, 17], and Reactome [18]. At the same time, advances in natural language processing (NLP) [19–24] have enabled systematic extraction of structured knowledge from the biomedical literature, such as the Semantic MEDLINE [25] Database (SemMedDB) [26]. Community-driven ontology development [27–31], literature curation, and the use of NLP together have driven growth of structured biomedical knowledge-bases, albeit in forms that are not semantically interoperable due to the use of different systems of concept identifiers, semantic types, and relationship types [32].

There have been numerous efforts to address the lack of semantic interoperability of structured biomedical knowledge, particularly in knowledge representation [33]. BIOZON [34], BioGraphDB [35], Hetionet [36], SPOKE [37, 38], EpiGraphDB [39], and DRKG [40] used standard sets of identifier types; and Bio2RDF [41], KaBOB [42], and HKGB [43] used ontologies [30, 31] for knowledge linking. ROBOKOP [44, 45], BioThings [46], and mediKanren [47, 48] use concept and relationship types from the recently-developed Biolink model [49–51]. Biolink is a high-level ontology that provides mappings of semantic types and relation types to other ontologies. Biolink advanced the field by (i) providing mappings of semantic types and relation types to other ontologies; (ii) standardizing and ranking preferred identifier types for various biological entities; and (iii) providing hierarchies of relation types and concept types needed to provide a semantic layer for biological knowledge graphs. In 2016, the National Center for Advancing Translational Sciences (NCATS) launched the Biomedical Data Translator project [52], a multi-institution effort to develop a distributed computational reasoning and knowledge exploration system for translational science. After a feasibility assessment phase in 2017–2020, the project began construction of the Translator system’s components such as reasoning agents, knowledge providers, and central controller system in 2020. The RTX-KG2 system is a registered knowledge provider within Translator. To ensure that Translator’s various systems can interoperate, Biolink has been adapted as the semantic layer for concepts and relations for knowledge representation within the Translator project.

Because biomedical knowledge-bases collectively use various semantically overlapping controlled vocabularies for concept types like diseases, drugs, phenotypes, and pathways, integrating knowledge into a graph entails grappling with the problem of multiple identifiers for a single concept; for example, the concept “paracetamol” has many identifiers, such as `UMLS:C0000970`, `DRUGBANK:DB00316`, `CHEBI:46195`, and `CHEMBL112`. While most biomedical knowledge graph efforts map concepts to canonical identifiers from semantic type-specific controlled vocabularies during initial graph construction, Monarch [53–55] constructed a linked graph of concept identifiers and then used clique detection to identify identical concepts before selecting a representative canonical identifier (a step that is called graph “canonicalization” [56]) for each clique. To date, biomedical knowledge graphs of which we are aware (with the exception of Bio2RDF [41]) are either canonicalized or standardized on a single identifier type for each semantic type, rather than providing *both* canonicalized and pre-canonicalized graphs; the latter form is important in order to support users that wish to apply their own canonicalization algorithm.

Previous efforts to develop integrated biomedical knowledge systems have used various database types, architectural patterns, and automation frameworks. For persistence, knowledge systems have used relational databases [34], distributed graph databases [33, 57], multimodal NoSQL databases [35, 57], RDF triple-stores [41, 42, 58], document-oriented databases [32, 46, 54], and—with increasing frequency [36, 37, 39, 44, 54]—the open-source graph database Neo4j ([github:neo4j/neo4j](https://github.com/neo4j/neo4j)). Knowledge systems have also differed in terms of the ingestion method used in their construction; many systems [32, 35, 41, 42, 54] utilized an extract-transform-load (ETL) approach, whereas others [44, 46, 59] used API endpoints to query upstream knowledge sources; one [39] blended both ETL and API approaches for knowledge graph construction. For automation, previous efforts have used general-purpose scripting languages [36, 37, 41, 42, 44, 59, 60], batch frameworks [32], rule-based build frameworks [33, 35, 61], semantic web-compliant build frameworks such as PheKnowLator [62], and parallel-capable systems such as Snakemake [63]. While previous efforts have resulted in biomedical knowledge graphs incorporating (individually) UMLS, SemMedDB, multiple major drug knowledge bases (such as ChEMBL and DrugBank), a standards-compliant semantic layer, and a parallel build system, so far as we are aware, none have incorporated all of these features in a single system providing both canonicalized and pre-canonicalized knowledege graphs.

## Introduction

We have developed RTX-KG2, an open-source biomedical knowledge graph representing biomedical concepts and their relationships. RTX-KG2 integrates 70 sources including the major sources UMLS, SemMedDB, ChEMBL, DrugBank, SMPDB, Reactome, KEGG, and UniProtKB using a modular build system leveraging the parallel-capable workflow framework, Snakemake. The semantic layer for RTX-KG2 is based on the standards-based Biolink model and it is provided in two stages, a pre-canonicalized graph version (RTX-KG2pre, in which semantically duplicated concepts with distinct identifiers are distinct nodes) and a canonicalized version (RTX-KG2c) in which equivalent concepts described using different identifier systems are identified as a single node. These key design choices reflect the goals of (i) supporting interoperability and composability with other biomedical knowledge-bases in the Translator system and (ii) providing a comprehensive knowledge graph with a standards-based semantic layer that is amenable to computational reasoning. Both RTX-KG2pre and RTX-KG2c are directed multigraphs with node and edge annotations that follow the Biolink model. The software repository for RTX-KG2, including all code to build the database, is publicly available at the [github:RTXteam/RTX-KG2](https://github.com/RTXteam/RTX-KG2) GitHub project. Users can access RTX-KG2 content via any of three channels: (i) a single-file download version of the canonicalized RTX-KG2 knowledge graph (KG2c) (or, if needed, the pre-canonicalized RTX-KG2pre knowledge graph); (ii) a publicly accessible, SmartAPI [64]-registered API for querying RTX-KG2; and (iii) a web browser interface for querying RTX-KG2. RTX-KG2 uses an ETL approach for knowledge graph construction and it automates builds using Snakemake; together, these enable efficient knowledge graph construction. RTX-KG2 is a built-in knowledge database for ARAX (Autonomous Relay Agent X) [65], a Web-based computational biomedical reasoning system that our team is also developing for answering translational science questions such as questions related to drug repositioning, identifying new therapeutic targets, and understanding drug mechanisms-of-action. We are developing RTX-KG2 and ARAX as a part of the NCATS Translator project. Here, we enumerate the knowledge sources that are incorporated into RTX-KG2 (Sec. 2.1); outline the processes for building RTX-KG2pre from its upstream knowledge sources (Sec. A.1) and for building the canonicalized RTX-KG2c (Sec. 2.2); describe the schema for RTX-KG2 (Sec. 2.3); describe the RTX-KG2 build system software (Sec. A.3); provide statistics about the size and semantic breadth of RTX-KG2 (Sec. 2.5); and discuss how it is being used for translational reasoning as well as in conjunction with the ARAX system (Sec. 3).

## 2 Construction and Content

In this section, we describe how RTX-KG2 is constructed; provide an overview of its graph database schema; and summarize its content in terms of sources, semantic breadth, and size. The overall build process, along with the various outputs of RTX-KG2, is depicted in Figure 1. Broadly speaking, the RTX-KG2 build system does four things: it (i) loads information from source databases (blue triangles in Fig. 1) via the World Wide Web as described in Section 2.1); (ii) integrates the knowledge into a precursor knowledge graph called RTX-KG2pre (upper green hexagon in Fig. 1) and hosts it in a Neo4j database (upper orange cloud in Fig. 1) as described in Section A.1; (iii) coalesces equivalent concept nodes into a canonicalized knowledge graph called RTX-KG2c (brown circle in Fig. 1) as described in Section 2.2, with a schema that is described in Section 2.3; and (iv) provides various knowledge graph artifacts and services as described in Section 2.5. We provide technical details of the RTX-KG2 build system in Section A.3.

**Figure 1:**
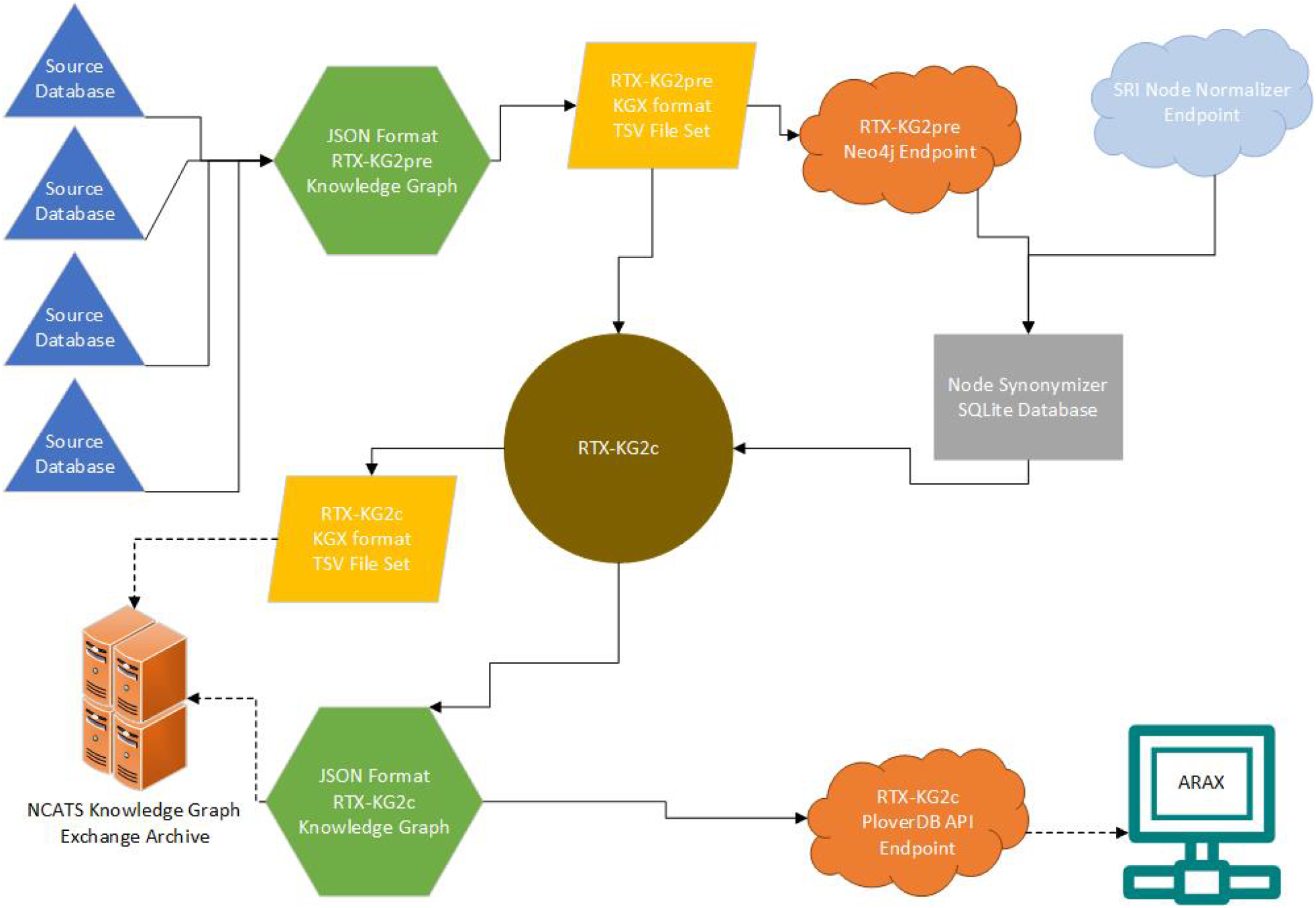
Overall Workflow of RTX-KG2. Blue triangle: individual external source; light blue cloud: external API endpoint; yellow parallelogram: tab-separated value (TSV) file-set; green hexagon: JavaScript Object Notation (JSON) File; orange cloud: API endpoint output; grey rectangle: SQLite [66] database; brown circle: abstract object-model representation of KG2c; turquoise computer: user/client computer; orange server: Translator knowledge graph exchange (KGE) server.

### 2.1 Sources and their file formats

RTX-KG2 integrates 70 sources (Table 1), 50 of them via a *resource description framework (RDF)-based* ingestion method and 20 of them via a *direct-to-JSON* ingestion method.

**Table 1:** RTX-KG2 integrates 70 knowledge sources into a single graph. Each row represents a server site from which sources were downloaded. Columns as follows: *Name*, the short name(s) of the knowledge sources obtained or the distribution name in the cases of UMLS and OBO Foundry; *#*, the number of individual sources or ontologies obtained from that server; *Format*, the file format used for ingestion (see below); *Method*, the ingestion method used for the source, either `D2J` for direct-to-JSON or `RBM` for the RDF-based method. File format codes: `CSV`, comma-separated value; `DAT`, SWISS-PROT-like DAT format; `JSON`, JavaScript object notation; `OWL`, OWL in RDF/XML [67] syntax; `RRF`, UMLS Rich Release Format [68]; `SQL`, structured query language (SQL) dump; `TSV`, tab-separated value; `XML`, extensible markup language. Other abbreviations: NCBI, National Center for Biotechnology Information; EMBL, European Molecular Biology Laboratory.

| Name | # | Description | Format | Method |
| --- | --- | --- | --- | --- |
| Biolink [49, 50] | 1 | Biolink Model (semantic layer) | TTL | RBM |
| ChEMBL [14, 69] | 1 | EMBL Chemogenomic Database | SQL | D2J |
| DGIdb [70] | 1 | Drug Gene Interaction Database | TSV | D2J |
| DisGeNET [71] | 1 | Disease-Gene Associations | TSV | D2J |
| DrugBank [13] | 1 | Pharmaceutical Knowledge Base | XML | D2J |
| DrugCentral [72] | 1 | Online drug Compendium | SQL | D2J |
| Ensembl Gene [73] | 1 | Ensembl Human Gene annotations | JSON | D2J |
| EFO [74] | 1 | Experimental Factor Ontology | OWL | RBM |
| GO [75, 76] | 1 | Gene Ontology annotations | TSV | D2J |
| HMDB [77–80] | 1 | Human Metabolite Database | XML | D2J |
| IntAct [81, 82] | 1 | IntAct Molecular Interaction Database | TSV | D2J |
| Jensen Lab Diseases [83] | 1 | Gene to Diseases Dataset | TSV | D2J |
| KEGG [11, 84, 85] | 1 | Kyoto Encyclopedia of Genes and Genomes | API | D2J |
| miRBase [86–90] | 1 | MicroRNAs Dataset | DAT | D2J |
| NCBI Gene [91] | 1 | NCBI Human Gene annotations | TSV | D2J |
| OBO Foundry | 21 | OBO Foundry Ontologies (Table S1) | OWL | RBM |
| Orphanet [92] | 1 | Orphanet Rare Disease Ontology | OWL | RBM |
| PathBank [93–95] | 1 | Wishart Lab Pathway Databases | XML | D2J |
| Reactome [96] | 1 | Pathway Database | SQL | D2J |
| SemMedDB [26] | 1 | Semantic MEDLINE Database | SQL | D2J |
| SMPDB [16, 17] | 1 | Small Molecule Pathway Database | CSV | D2J |
| UMLS [97] | 26 | Unified Medical Language System (Table 7) | TTL | RBM |
| UniChem [98] | 1 | EBI Small Molecule Cross-refs | TSV | D2J |
| UniProtKB [15] | 1 | UniProt Knowledge Base | DAT | D2J |
| Total | 70 |  |  |  |

Sources are loaded in a specific order controlled by a configuration file, with precedence applying to the assignment of Biolink categories to nodes.

#### 2.1.1 RDF-based sources

Of the 50 RDF-based sources, the system ingests 27 in Terse RDF Triple Language (TTL [99]) format and 23 as OWL ontologies in RDF/XML format [67] (which we abbreviate here as “OWL”). Of the 27 TTL sources, 26 are from the UMLS, obtained as described in Sec. A.3.3; the remaining source is a TTL representation of the Biolink model, which defines the semantic layer for RTX-KG2, including hierarchies of concept types and relation types (see Sec. 2.5). In addition to concept type and relation type hierarchies, the Biolink model provides equivalence mappings of the Biolink types to classes in other high-level ontologies (such as `biolink:Gene` being equivalent to `SIO:010035`) and of the Biolink concept types to prioritized lists of identifier types for the concept type^1^. Each knowledge source’s concepts are assigned Biolink concept semantic types—which are called “categories” in the Biolink model—and relationships are assigned Biolink relationship types at the time that the source is ingested. All but two of the 23 OWL-format sources are ontologies from the OBO Foundry [31]; the remaining two OWL-format sources are the Experimental Factor Ontology (EFO) [74] and Orphanet Rare Disease Ontology [100].

#### 2.1.2 Direct-to-JSON sources

With the direct-to-JSON method, sources are ingested in seven different file formats (Table 2.1). One source, KEGG, is queried via an API rather than using a flat file download, due to the lack of a download option for users that do not have a commercial license. For the 20 direct-to-JSON sources, the RTX-KG2 system has one ETL module for each source, with each module producing a source-specific JSON file in the RTX-KG2 JSON schema (Sec. 2.3) (in contrast, for the 50 RDF-based sources, the system has a single ETL module for ingesting all sources together). The RDF-based method merges all of the OWL and TTL sources (class-based), without flattening the ontologies (i.e., preserving `rdfs:subClassOf` relationships) and generates a single JSON file. The hybrid design of RTX-KG2 balances the benefits of modularity (where it is feasible in the direct-to-JSON method) with the need for a monolithic ingestion module for ontologies where inter-ontology axioms are needed for determining semantic types at the ETL stage [101].

**Table 2:** Node and edge counts for various knowledge graphs.

|  | Nodes | Edges |
| --- | --- | --- |
| HETIONET, v1 [36] | 47,031 | 2.3 million |
| SPOKE ver. 20190707 [38] | 2.15 million | 6.16 million |
| SRI Reference KG, ver. 2.0 | 20.2 million | 41.6 million |
| ROBOKOP [45] | 6 million | 140 million |
| RTX-KG2.7.3pre | 10.2 million | 54.0 million |
| RTX-KG2.7.3c | 6.4 million | 39.3 million |

### 2.2 Building RTX-KG2c, the canonicalized version of RTX-KG2

Because the various ontologies that RTX-KG2pre ingests often represent the same concept using multiple different identifiers, some of the nodes in RTX-KG2pre represent equivalent concepts. For example, *Parkinson’s disease* is represented by several nodes in RTX-KG2pre, such as the nodes with identifiers `MONDO:0005180`, `DOID:14330`, `EFO:0002508`, and `MESH:D010300`, many of which are connected in RTX-KG2pre with relationships of type `biolink:same as` or non-transitive generalizations of that relationship type. In our work on RTX-KG2pre, we found that coalescing nodes for semantically equivalent concepts into single nodes facilitates reasoning by reducing the complexity of graph paths that represent answers for common translational questions. Thus, to enhance the utility of RTX-KG2 for translational reasoning, we created a version of RTX-KG2 called RTX-KG2canonicalized (abbreviated in this work as RTX-KG2c) in which semantically equivalent nodes are coalesced to a single concept node. In brief, building RTX-KG2c from RTX-KG2pre is carried out in five steps:

1. RTX-KG2pre nodes and edges are loaded from the RTX-KG2pre TSV files;
2. the set of nodes is partitioned into disjoint subsets of equivalent nodes;
3. from each group of equivalent nodes, a canonical node identifier is chosen, added to RTX-KG2c, and annotated with the identifiers of its synonymous nodes (along with other information merged from the synonymous nodes);
4. edges from RTX-KG2pre are remapped to refer only to the canonical node identifiers; and
5. edges with the same subject, object, and Biolink predicate are merged.

For Steps 2–3, the RTX-KG2 build system uses the ARAX [65] system’s *Node Synonymizer* service (see Sec. A.1.1 for details). The RTX-KG2c graph is serialized in JSON format (see Sec. 2.3), archived in a GitHub large file storage (LFS) repository (see Sec. 6.3), and imported into a custom-built, open-source, in-memory graph database, PloverDB ([github:RTXteam/PloverDB](https://github.com/RTXteam/PloverDB)). The build process for RTX-KG2c is Python-based and has comparable hardware requirements to the RTX-KG2pre build process (see Sec. A.3.1). Formally, the approach used in building RTX-KG2c is concept-oriented as opposed to the realist methodology underlying OBO Foundry ontologies [102].

### 2.3 RTX-KG2 schema and RTX-KG2pre Biolink compliance

The RTX-KG2.7.3 knowledge graph follows the Biolink model (version 2.1.0) for its semantic layer and (in RTX-KG2pre) its schema. RTX-KG2 uses Biolink’s category hierarchy for its concept (node) types (Fig. 2) and Biolink’s predicate hierarchy for its relationship (edge) types (Fig. 3). When mapping terms from their original sources to the Biolink terminology, the RTX-KG2 build system consults the Biolink model’s internal mappings in order to detect any inconsistencies between the two. Because relationship terms that are highly specific are often mapped to more generalized terminology, the original source’s phrasing is preserved in the `relation` property^2^. In addition to mapping upstream source relations to Biolink predicates, the RTX-KG2 build process coalesces edges that have the same end nodes and the same predicate (it does, however, preserve the provenance information from both of the coalesced edges). The schema of the JSON serialization of RTX-KG2pre is documented in detail in the RTX-KG2 project area [github:RTXteam/RTX-KG2](https://github.com/RTXteam/RTX-KG2/blob/master/README.md#schema-of-the-json-kg2) and summarized in Sec. A.2. In brief, RTX-KG2 is serialized as a JSON object with keys `nodes` and `edges`, with the `nodes` object containing a list of serialized objects for the concept nodes in the graph, and with `edges` containing a list of serialized objects for the subject-object-relationship triples in the graph.

**Figure 2:**
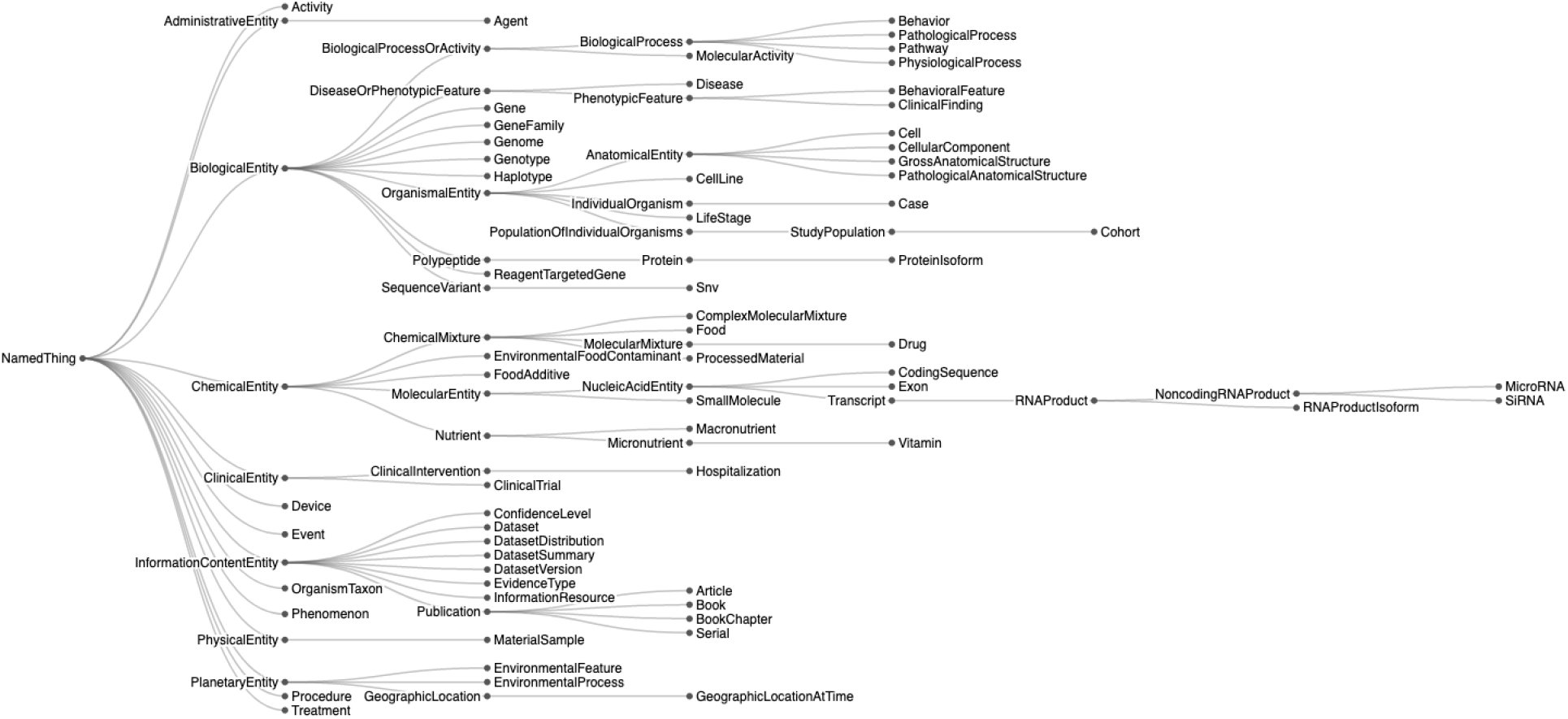
Node concept types in RTX-KG2.7.3 are based on the Biolink model version 2.1.0 [49, 50].

**Figure 3:**
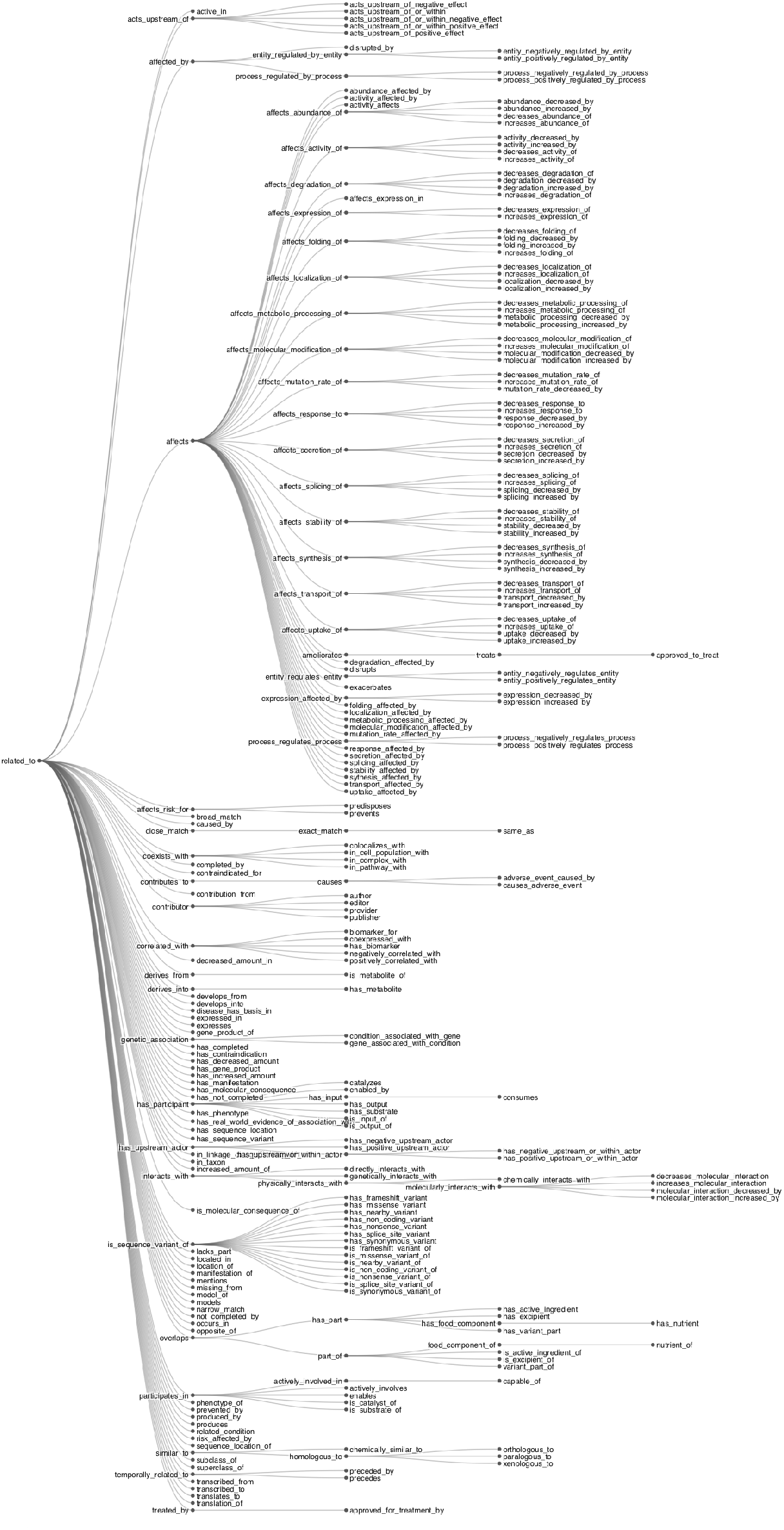
Edge predicate types in RTX-KG2.7.3 are based on the Biolink model version 2.1.0.

### 2.4 Quality control and reproducibility

The RTX-KG2 build process incorporates multiple layers of quality control, including both automated and manual procedures (see Sec. A.1 for details). As the first step in the build process, scripts validate the consistency of the RTX-KG2 semantic layer with the Biolink model. During knowledge integration, relationships whose subject or object nodes are not present in the knowledge graph are logged for offline investigation. A report of statistics on the RTX-KG2 knowledge graph is generated—including (i) node counts by knowledge source and by semantic type and (ii) edge counts by source and by relationship type—both before and after redundant edges are joined in the merge process. The procedure for RTX-KG2 builds includes a script-facilitated comparison of that report for the new build with the equivalent report for the previous build, in order to enable the build supervisor to recognize anomalously large (e.g., more than three-fold) changes in node or edge counts conditioned on source, category, or predicate. Once a new RTX-KG2c build is installed into the RTX-KG2 query API, functional correctness of the live API is verified using a Python-based test suite. Reproducibility of the RTX-KG2 build is enhanced by the intentional choice of using an ETL approach based on flat-file exports knowledge-source databases. To aid with versioning, most sources have their version stored in the name attribute in the node in RTX-KG2 that represents the source database.

### 2.5 RTX-KG2 content and statistics

The latest released version of RTX-KG2pre as of this writing, RTX-KG2.7.3, contains 10.2 million nodes and 54.0 million edges. Each edge is labeled with one of 77 distinct predicates (Biolink relationship types) and each node with one of 56 distinct categories (Biolink concept semantic types). In terms of frequency distribution, there is over six decades of variation across node categories (Fig. 4) and edge predicates (Fig. 5), with the dominant category being `OrganismTaxon` (reflecting the significant size of the NCBI organism classification ontology [103]) and the dominant predicate being `has_participant` (reflecting the significant size of the PathBank database [93]). Figure 5 shows a breakdown of edges in KG2.7.3 by their Biolink predicate. KG2.7.3c contains 6.4 million nodes and 39.3 million edges, which is approximately 62% of the nodes and 73% of the edges of KG2.7.3pre, reflecting an expected reduction in node count due to canonicalization as well as due to post-canonicalization edge merging. By number of edges, the largest contributing knowledge source for RTX-KG2.7.3pre is SemMedDB, which has 19.3M edges (comprising about a third of the edges in the graph), followed by PathWhiz (13.7M edges), NCBI Taxonomy (3.6M edges), and DrugBank (2.8M edges).

**Figure 4:**
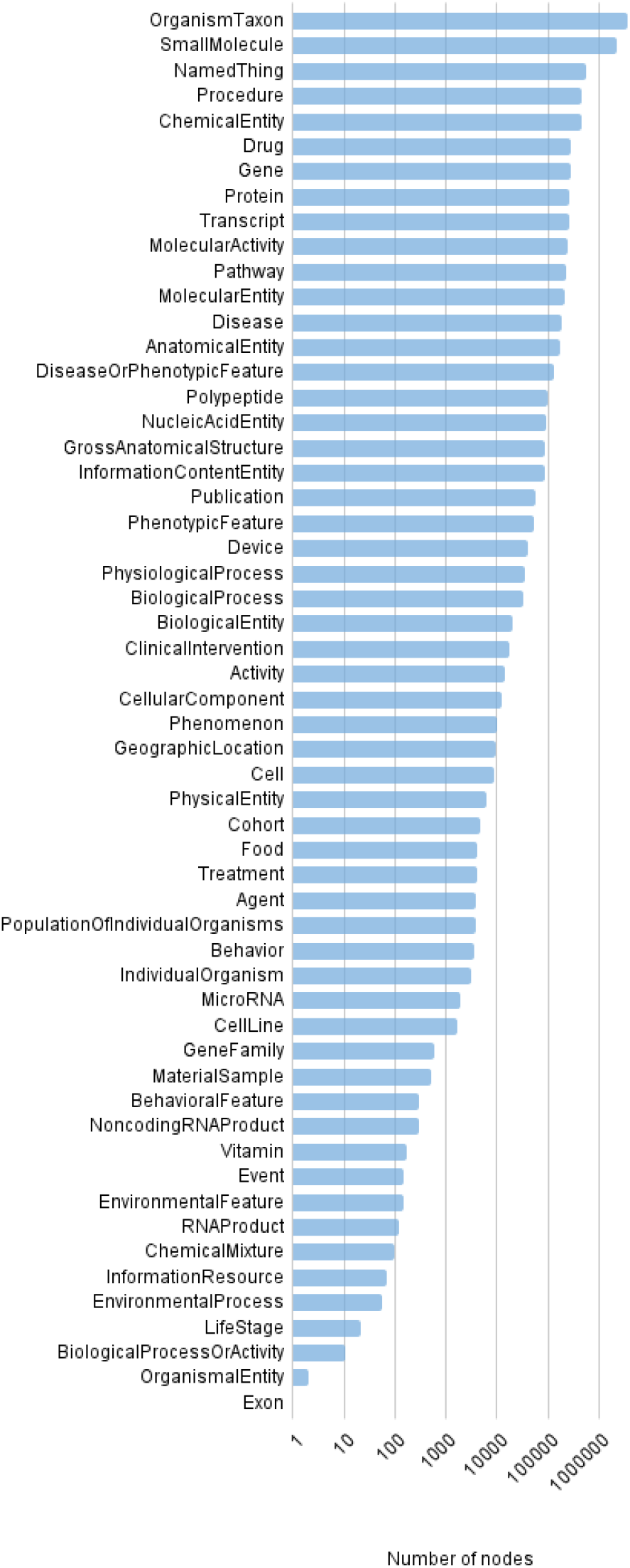
Number of nodes in RTX-KG2.7.3pre, by category.

**Figure 5:**
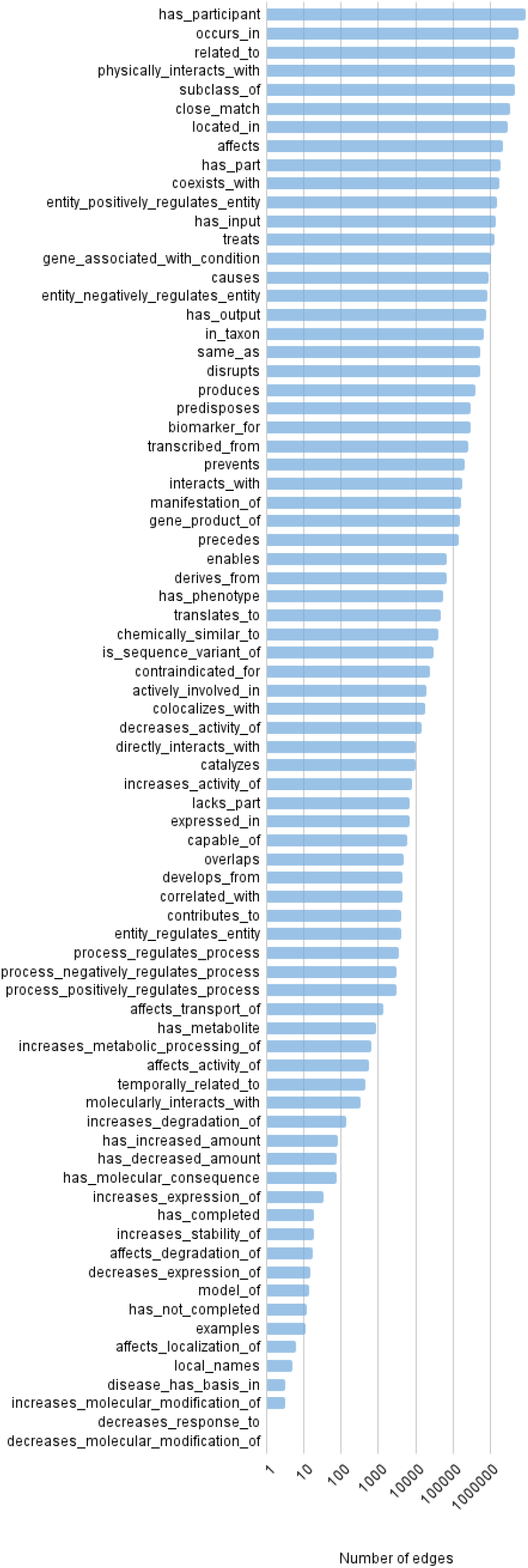
Number of edges in RTX-KG2.7.3pre, by predicate.

In terms of their total (i.e., in+out) vertex degree distributions, both KG2pre and KG2c appear to be approximately scale-free (Fig. 7) with a power law exponent of 2.43, meaning that the frequency of concepts with connectivity *k* decreases as ∼*k*^*−*2.43^. Figure 6 highlights the frequencies of various combinations of subject node category and object node category appearing together in edges in KG2c, indicating (1) high levels of cross-category axioms among “molecular entity”, “small molecule”, and “chemical entity” and (2) high levels of connections between “pathway” and “molecular entity”, “small molecule”, “molecular activity”, “organism taxon”, “anatomical entity”, and “transcript”. Note that the category-category frequency heatmap is not expected to be symmetric for a knowledge graph (such as RTX-KG2) with a high proportion of relationship types that have non-reflexive subject-object semantics.

**Figure 6:**
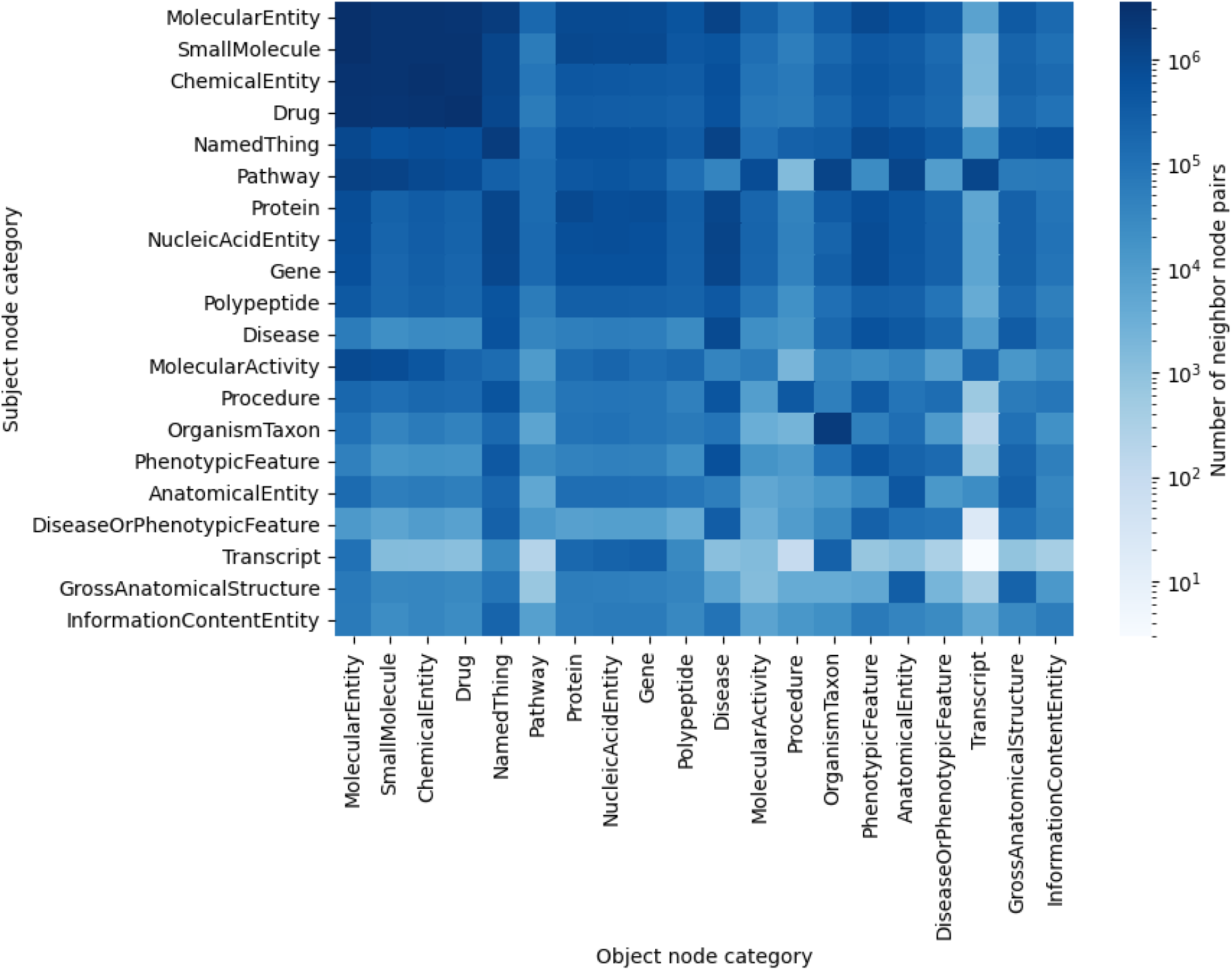
Node neighbor counts by category for the top 20 most common categories in RTX-KG2.7.3c. Each cell captures the number of distinct pairs of neighbors with the specified subject and object categories.

**Figure 7:**
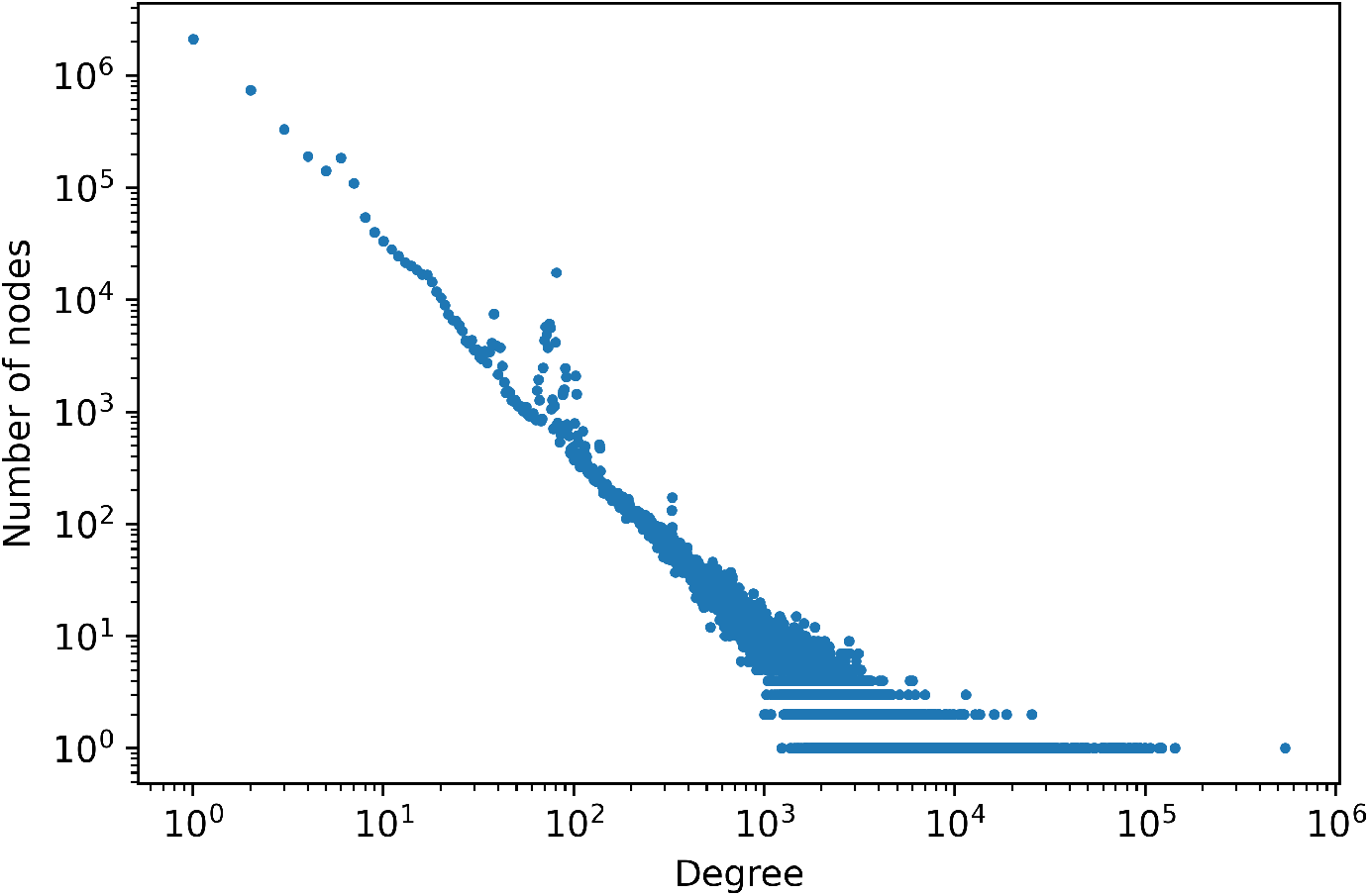
Node degree (in+out) distribution of RTX-KG2.7.3c.

### 2.6 RTX-KG2 access channels

The complete software code for building RTX-KG2 and for hosting an indexed RTX-KG2 graph database in Neo4j is publicly available in an open-source repository (see Sec. 6.3). In addition, the content of the latest RTX-KG2 graphs (version KG2.7.3) that we have built can be easily accessed via three different channels (see Sec. 6.3), depending on the use-case: (i) serialized flat-file download (as described below); (ii) REpresentational State Transfer (REST) [104] API (i.e., a web API); or (iii) web browser user interface, through the ARAX system. Tab-separated value (TSV) serializations of RTX-KG2pre and JSON serializations of RTX-KG2c are available in a public GitHub repository via the `git-lfs` file hosting mechanism, and their schemas are documented as described in Sec. 2.3 and in the RTX-KG2 documentation sections that are linked therein. RTX-KG2c can be queried via a REST API that implements the open-standard Translator Reasoner API (TRAPI) specification ([github:NCATSTranslator/ReasonerAPI](https://github.com/NCATSTranslator/ReasonerAPI)) and that is registered via the SmartAPI [64] framework and therefore discoverable using SmartAPI-associated tooling such as BioThings Explorer [46]. The RTX-KG2 API enables one-hop querying of the knowledge graph; queries are internally serviced by the PloverDB in-memory graph database (see Sec. 2.2). The ARAX API (arax.rtx.ai/api/arax/v1.2/openapi.json), which itself queries the RTX-KG2 API, can be used to achieve multi-hop RTX-KG2 queries. Further, RTX-KG2c is archived in Biolink Knowledge Graph eXchange [49] TSV format (KGX TSV format, documented at [github:biolink/kgx](https://github.com/biolink/kgx)) through the Translator Knowledge Graph Exchange (KGE; see Figure 1) archive and registry system ([github:NCATSTranslator/Knowledge_Graph_Exchange_Registry](https://github.com/NCATSTranslator/Knowledge_Graph_Exchange_Registry)) (currently in testing phase).

## 3 Utility and Discussion

### 3.1 Uptake and adoption

Due to its comprehensiveness and/or its speed, RTX-KG2 is already being used as a core knowledge provider (see [github:NCATSTranslator/Translator-All/wiki/KG2](https://github.com/NCATSTranslator/Translator-All/wiki/KG2)) or knowledge graph by five diverse reasoning agents within the Translator system: ARAX [65], which our team developed and which provides sophisticated workflow capabilities and overlay of virtual edges for associations based on literature co-occurrence or network structural equivalence; mediKanren, which provides sophisticated network motif-finding and path-finding using the miniKanren logic programming language; BioThings Explorer, the engine for autonomous querying of distributed biomedical knowledge, described in Section 1; ARAGORN ([github:ranking-agent/aragorn](https://github.com/ranking-agent/aragorn)), a reasoning agent that has unique capabilities for coalescing and ranking knowledge subgraphs; and the Explanatory Agent ([github:NCATSTranslator/Explanatory-Agent](https://github.com/NCATSTranslator/Explanatory-Agent)), a reasoning agent that uses natural language-understanding models in order to explain and rank results.

Use of the RTX-KG2 API appears to be increasing over time, with an average of 1,084 queries per day over the 9 months prior to this writing (September 2021 - June 2022) vs. an average of 1,417 queries per day over the last 3 months (April - June 2022), a 1.3-fold increase. Programs that query RTX-KG2 may optionally identify themselves; of the 51% of queries from the last 9 months in which the submitter was identified, approximately 63% were by the Explanatory Agent, 23% by ARAX, 7% by BioThings Explorer, and 7% by ARAGORN. Rather than using the RTX-KG2 API, the mediKanren reasoning agent uses a bulk download of RTX-KG2pre in conjunction with their own canonicalization algorithm.

In addition to its primary intended use-case for on-demand knowledge exploration and concept-specific reasoning, the RTX-KG2 knowledge graph can be used as a structure prior for data-driven network inference, for example, causal network learning. We have recently described a computational method, *Kg2Causal* [105], for using a general-purpose biomedical knowledge graph to extract a network structure prior distribution for data-driven causal network inference from multivariate observations. Using the predecessor graph, RTX-KG1 [59], we found that using a general knowledge graph as a prior significantly improved the accuracy of data-driven causal network inference compared to using any of several uninformative network structure priors [105]. To the extent that it incorporates multiple graph structural variations, RTX-KG2 can also be used as a test-bed for evaluating the performance of structurally generalizable graph analysis methods such as a subset of us have done for the case of a structurally generalizable node-node similarity measure [106].

Another application of RTX-KG2 is as training data for drug repurposing models; previous work by our team utilized RTX-KG2’s predecessor, RTX-KG1, to train a random forest model that predicts novel drug treatments for diseases [107]. More recent extensions to this work have utilized the canonicalized version of RTX-KG2 as training data.

### 3.2 Comparison to other knowledge graphs/providers

To objectively evaluate the size and semantic richness of RTX-KG2 in comparison to other biomedical knowledge graphs, we compared it to four other knowledge graphs that are in active use for translational applications: Hetionet [36], SPOKE [37, 38], the SRI Reference Knowledge Graph [108], and ROBOKOP [44, 45]. For the counts of meta-triples (i.e., counts of edges with a given pattern of subject category, object category, and predicate, which provide quantitative information about the richness of the knowledge graph for providing relationships of particular types), we accessed the ROBOKOP and SPOKE graphs via their SmartAPI-registered Translator API (“TRAPI”) endpoints on March 15, 2022. For Hetionet and ROBOKOP, we used published node and edge counts [36, 45]. For the SRI Reference Knowledge Graph, we downloaded the graph in KGX TSV format from the Biomedical Data Translator Knowledge Graph Exchange (KGE) and analyzed the graph locally.

Node and edge counts for various knowledge graphs are shown in Table 2. The large number of edges in the ROBOKOP knowledge graph versus RTX-KG2 reflects the latter’s practice of joining edges that have identical subject node, object node, and predicate type. The large number of nodes in the SRI Reference Knowledge Graph [108] versus RTX-KG2pre is largely due to more `InformationContentEntity` nodes (3.7 million vs. 144,396) and `SequenceVariant` nodes (2.4 million vs. 0) in the former.

Notably, RTX-KG2 contains a richer set of meta-triples (distinct combinations of subject node category, edge predicate, and object node category) versus the other knowledge graphs (Table 3); RTX-KG2c contains 4.6-fold more meta-triples than the second-ranked knowledge graph and 18.5-fold more meta-triples than the third-ranked knowledge graph, by meta-triple count. In general we would expect a graph with a greater number of meta-triples to be able to provide answers to a wider variety of queries, which is somewhat corroborated by the findings in the following paragraph. The finding that canonicalization increases the number of meta-triples can be understood as follows: since each node in RTX-KG2c has multiple categories, the number of meta-triples increases by the product of the count of subject node categories and the count of object node categories, for all subject-object node pairs joined by edges in the graph. The approximately 2.4-fold increase in the number of predicates in ROBOKOP versus in RTX-KG2 partially reflects the design choice in RTX-KG2 to standardize predicate directions and to orient triples so that for any inverted pair of predicates (e.g., “has part” and “part of”), only one of the predicates is used in the graph. Here, we have opted to compare RTX-KG2 to biomedical knowledge graphs that are canonicalized and still being updated (which excludes the pre-canonicalized Bio2RDF [41], which has not been updated since 2014).

**Table 3:** Numbers of unique node categories, edge predicates, and meta-triples for various knowledge graphs.

|  | Categories | Predicates | Meta-triples |
| --- | --- | --- | --- |
| SPOKE, TRAPI v1.2.0 API | 14 | 24 | 44 |
| SRI Reference KG, ver. 2.0 | 62 | 59 | 2,047 |
| ROBOKOP, TRAPI v1.2.0 API | 20 | 185 | 2,234 |
| RTX-KG2.7.3pre | 56 | 77 | 10,269 |
| RTX-KG2.7.3c | 56 | 77 | 41,225 |

To estimate the novelty of knowledge that RTX-KG2 provides over other knowledge providers, we ran a diverse set of one-hop queries^3^ through the ARAX reasoning agent and measured how many results were returned when ARAX was allowed vs. was not allowed to use RTX-KG2 as one of its knowledge providers. Omitting RTX-KG2 and relying on its 12 other Translator knowledge providers resulted in an average of 46% of the results compared to when RTX-KG2 was included. Figure 8 details the results for each query tested.

**Figure 8:**
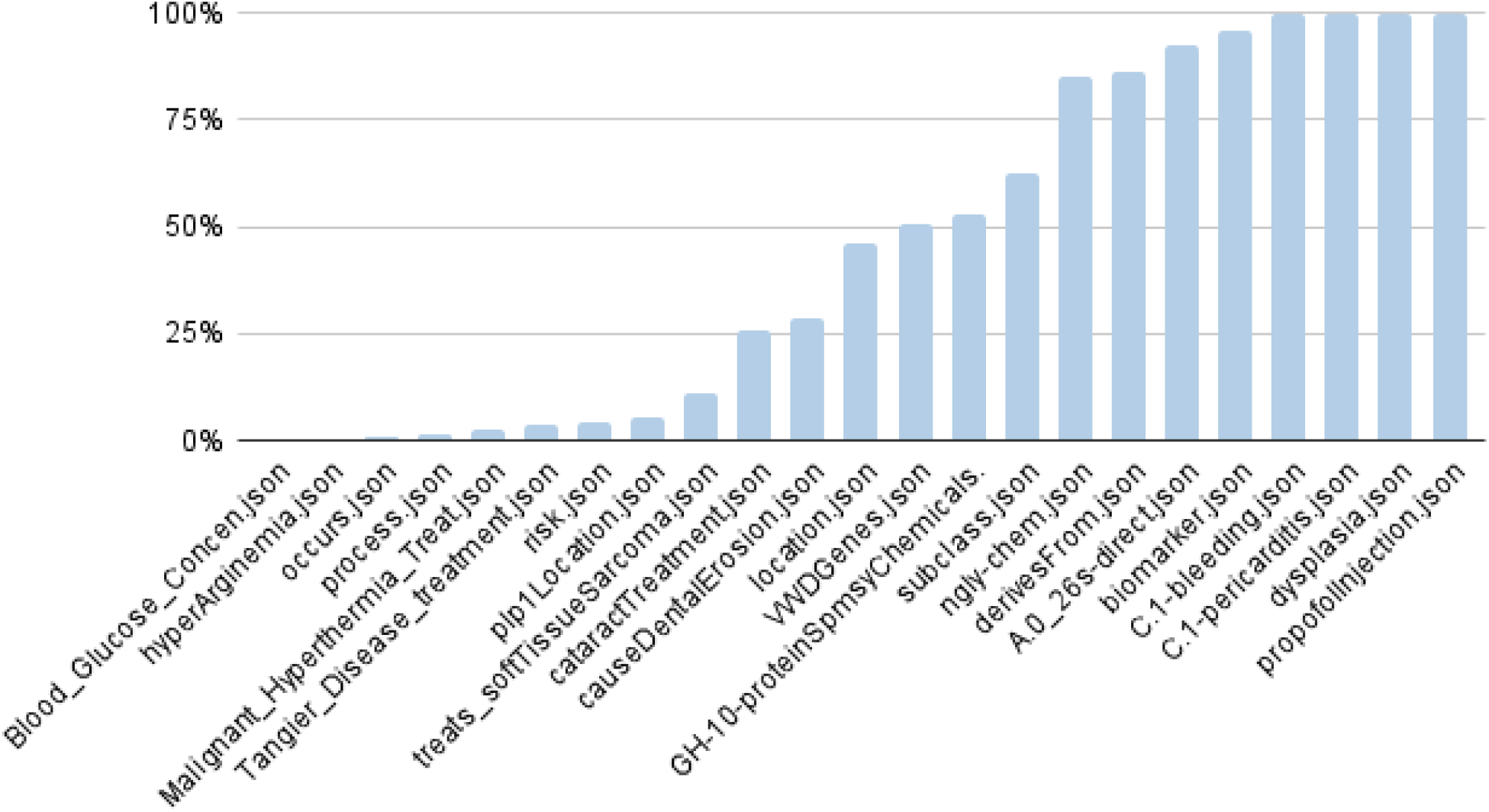
The proportion of results ARAX obtains for various one-hop queries when it is *not* allowed to use RTX-KG2 as one of its knowledge providers vs. when it is allowed to use RTX-KG2. A result of 100% means that KG2 provided no additional answers over ARAX’s other 12 Translator knowledge providers for that query; 0% means that all of ARAX’s results for that query came from RTX-KG2.

### 3.3 Example use-cases

To help illustrate how to use RTX-KG2, an example RTX-KG2 API query is provided below. This query (which is written in TRAPI format) asks RTX-KG2 for genes related to Adams-Oliver syndrome:

~~~
curl -X ‘POST’ \
‘https://arax.rtx.ai/api/rtxkg2/v1.2/query‘ \
-H ‘accept: application/json’ \
-H ‘Content-Type: application/json’ \
-d ‘{
“message”:{
   “query_graph”:{
       “edges”:{
           “e00”:{
               “subject”:”n00”,
               “object”:”n01”,
               “predicates”:[
                  “biolink:related_to”
               ]
           }
       },
       “nodes”:{
          “n00”:{
              “ids”:[
                 “MONDO:0007034”
              ],
              “is_set”:true
          },
          “n01”:{
             “categories”:[
                “biolink:Gene”
             ]
           }
         }
       }
     }
}’
~~~

The response that RTX-KG2 returns to this query is also in TRAPI format and contains a ranked list of `results` and a `knowledge_graph`, per the TRAPI specification. The structure of each result object matches that of the `query_graph`, while the `knowledge_graph` contains all of the nodes and edges used in the results, decorated with evidence, provenance, and other information. As of June 30, 2022, this query returns 41 results, the top 10 of which are the genes *DOCK6, DLL4, NOTCH1, EOGT, RBPJ, ARHGAP31, CDC42, OGT, LFNG*, and *PAMR1*.

As stated in Sec. 2.6, our ARAX reasoning system [65] provides a web browser interface (which is publicly available as described in Sec. 6.3) that can be used to both construct queries of RTX-KG2 and browse ranked results from those queries. The browser interface provides various capabilities including node synonymization; graphical layout and exploration of annotated result-graphs; and a graphical, point-and-click query builder. RTX-KG2pre serves as the data foundation for the ARAX Node Synonymizer (described in Section A.1.1), which is accessible via the ARAX UI (example: arax.rtx.ai/?term=naproxen) or programmatically via the ARAX API:

~~~
curl -X ‘GET’ \
  ‘https://arax.rtx.ai/api/arax/v1.2/entity?q=naproxen‘ \
  -H ‘accept: application/json’
~~~

BioThings Explorer also provides a web browser-based user interface (biothings.io/explorer/query) including a query graph builder that can be used to query RTX-KG2 among other reasoning agents and knowledge providers. Figure 9 shows the example Adams-Oliver query in the BioThings Explorer query graph builder; click the “Query ARS” button to run the query and then the “Open ARS” button to go to a different user interface (provided by ARAX), in which users can select and explore result sets for different reasoning agents, some of which use RTX-KG2 as a knowledge provider (ARAX, BTE, ARAGORN, Explanatory Agent, and Unsecret Agent (which is based on mediKanren)).

**Figure 9:**
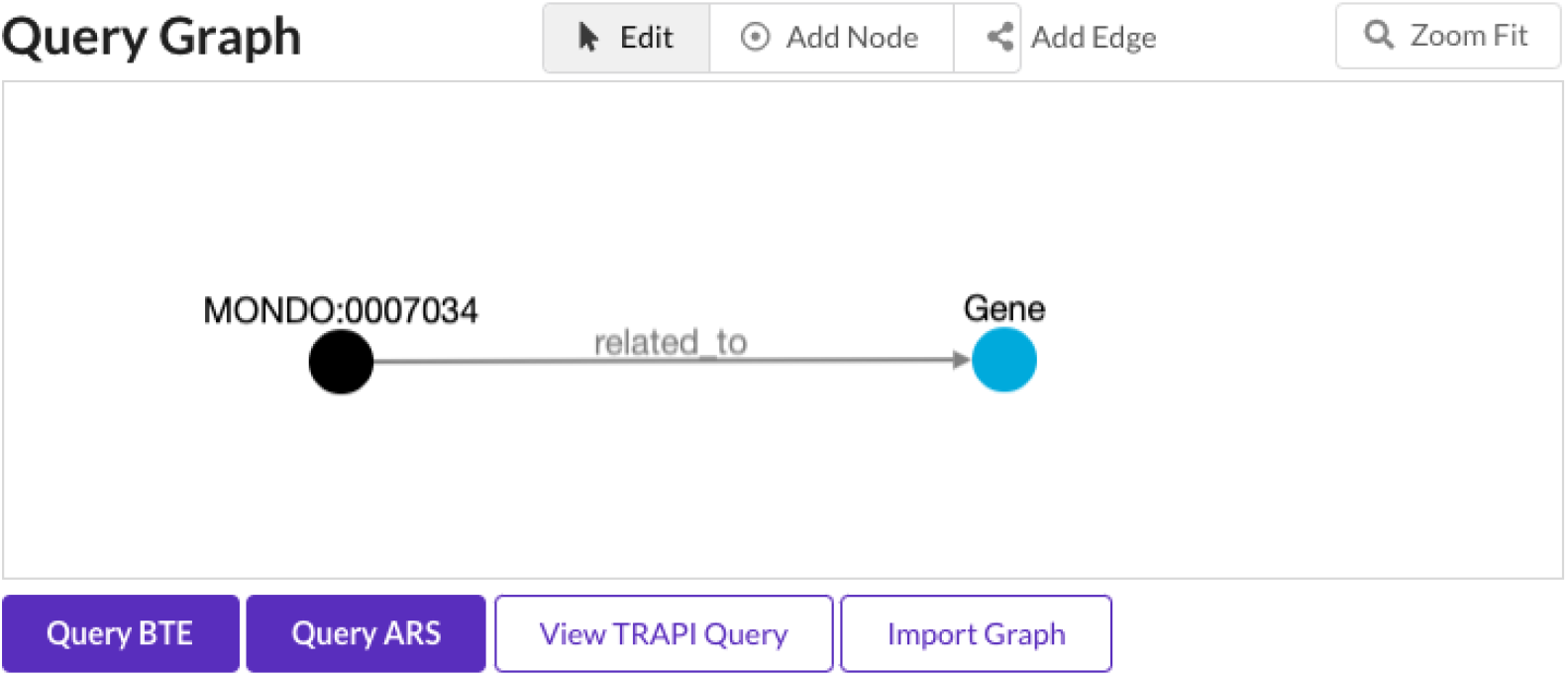
The BioThings Explorer query graph builder, which can be used to query RTX-KG2 among other Translator reasoning agents and knowledge providers.

### 3.4 Discussion

In designing RTX-KG2, we chose five design principles that guided our selection of knowledge sources to incorporate as well as the architecture of the RTX-KG2 build system:

1. Source is publicly available in a flat-file (e.g., TSV, XML, JSON, DAT, or SQL dump) that can be downloaded via a script
2. Source is being maintained and updated periodically
3. Source provides knowledge triples that complement (i.e., not duplicate) what is already in RTX-KG2
4. Source connects concept identifier types that are already in RTX-KG2
5. Ideally, source provides knowledge based on human curation

Principle 1, and the deliberate choice of using an ETL approach, theoretically would allow RTX-KG2 to be reconstructed consistently and independently of the state of external APIs^4^. This is useful for reproducibility, since each knowledge source is stored in its original downloaded form as a build artifact. Using flat files instead of API interfaces also increases the probability that a future build can be completed successfully at any time, since it does not rely on multiple web services to be up for an extended period of time. Additionally, it is in many (though by no means all) cases computationally faster to ETL a file than to dynamically query an API over the Internet. With the ETL approach, for inter-ontology axioms, full interoperability is required and thus, generally full resource import (versus partial import as proposed previously [109]) is used. Development of RTX-KG2 is ongoing and our team welcomes recommendations of new sources to include, via issue reports on the RTX-KG2 GitHub project page (see Sec. 6.3).

In selecting the 70 sources for RTX-KG2, we generally adhered to the aforementioned principles but made a few exceptions based on specific trade-offs. For Principle 1, for one source (as described in Sec. 2), we used an API rather than a file download, and for the “via a script” part of Principle 1, we manually downloaded source dump files for DrugBank, UMLS, and SemMedDB (due to the download page requiring a login using a web browser) and RepoDB (due to its information on drug approval status). Some large databases such as SNOMED CT and MedDRA were not included in the UMLS ETL because they have additional restrictions under the UMLS Metathesaurus License (Appendix 2 and Section 12.3, respectively). For Principle 2, an exception was miRbase, due to the lack of a clear alternative source. For Principle 3, partial exceptions were made for the various pathway databases such as Reactome, PathWhiz/SMPDB, and KEGG, which have many overlapping pathways but which also had systems of pathway identifiers that needed to be included in RTX-KG2. Further, each of the pathway databases has different strengths: PathWhiz/SMPDB offer useful links to HMDB and DrugBank; Reactome is popular, trusted, and is well connected with sources like GO and CHEBI; and KEGG CURIEs are popular with users and link to CHEMBL, CHEBI, and GO. The primary exception to Principle 5 is SemMedDB which is based on natural-language processing of biomedical research article abstracts to extract knowledge triples. SemMedDB is particularly useful for downstream reasoning because of its breadth across biomedical literature and because it includes source article references for each triple. However, in the NLP arena, new methods such as REACH [21] and EIDOS [22] have been proposed that promise improved accuracy for determining event polarity (REACH) and detecting causal mechanisms (EIDOS); the potential of these methods to be applied to the full biomedical literature corpus remains to be fully explored.

It is a challenge to balance the importance of manually curated knowledge resources with those that provide numerical data and provenance (such as supporting publications) of their assertions. While these two are not mutually exclusive *per se*, relatively few knowledge sources seem to provide both. Increasingly, reasoning agents in the Translator system will use structured provenance and confidence information/annotations for edges in knowledge graphs such as RTX-KG2; the issue of knowledge sources that are important connectors in translational reasoning but do not provide structured provenance information is an ongoing problem [110]. A second notable challenge for computational biomedical reasoning is that of conflicting information between sources, which can occur due to data entry error at upstream sources, updating of concept identifiers or identifier prefixes, or changes in the semantic layer. In our experience, careful scrutiny of the build report (described in Sec. 2.4) is essential to catch systemic problems so that they can be addressed before the build is put into production. On the other hand, for ameliorating the effects of localized/incidental issues due to random errors in curated upstream sources (or incorrect triples called by the NLP algorithm used in SemMedDB), we have found that in computational reasoning, overlaying of multiple sources of quantitative evidence of association (such as co-occurrence in the literature or graph structural similarity) to be beneficial within the ARAX system [65]. Another source of potential errors is due to the absence of contextual information for a triple, for example, an interaction that applies only in a specific anatomic context; work is ongoing to extend the Biolink model and RTX-KG2 with qualifier semantics for annotating core triples with such contextual information, as described below.

Use of Biolink for the semantic layer for RTX-KG2 provides advantages both within and outside the Translator ecosystem. Normalization of the more than 1,228 source relationship types to a smaller set of predicates is necessary in order to produce a knowledge graph that is amenable to rule-based reasoning. The choice of 77 predicate types in Biolink is a trade-off, balancing simplicity of reasoning with semantic precision. Work on the Biolink model is ongoing to eliminate this trade-off by allowing for annotation of triples with statement qualifiers, subject qualifiers, object qualifiers, and typed associations. Further, substantial tooling within the Biolink project [51] has been developed to (i) provide ports of the Biolink model to six different open-standards representations (e.g., OWL, TTL, ShEx, JSON-schema, and GraphQL), (ii) provide a turn-key software package (KGX [111]) for import, export, and validation of Biolink-based knowledge graphs in various formats (e.g., TSV and JSON), (iii) provide a turn-key software library for programmatic manipulation of the Biolink model as Python or Java classes, and (iv) provide comprehensive cross-ontology mappings between Biolink and other high-level ontologies, for both entity types and predicates. Collectively, these efforts are expected to enable computational biomedical reasoning efforts that are outside Translator to interoperate at the semantic layer in order to integrate or query structured knowledge from RTX-KG2. The design of RTX-KG2 further prioritizes reproducibility by release-tagging, documenting versions of upstream sources (and the Biolink model version) with the RTX-KG2 release, including Biolink schema validation as a part of the release process, and having the build system environment setup automated and under source code control.

Our observation that RTX-KG2c has a scale-free degree distribution is consistent with previous reports from empirical studies of text-based semantic networks [112] and ontologies [113]. Further, the subgraph of RTX-KG2 that arises from the SemMedDB NLP analysis of PubMed/MEDLINE [114] and from the Gene Ontology [115] would be expected to follow a scale-free degree distribution, in accordance with Zipf’s Law. Quantitatively understanding the degree distribution of the knowledge graph (whether conditioned on the knowledge source, the relationship type, or the participating entity types) is relevant to quantitatively modeling the epistemic value of an edge in the graph (consider, for example, the downweighting by neighboring vertex out-degree in the PageRank algorithm [116]).

As described in Sec. 1, the Neo4j graph database management system is used by many biomedical knowledge graphs including Hetionet, Monarch, SPOKE, EpiGraphDB, and ROBOKOP. Using Neo4j to host RTX-KG2 has both benefits (specifically, the flexibility of the Cypher query language [117]) and drawbacks (in our testing, slow JSON loading performance and slow response times in comparison to an in-memory database). It was due to its drawbacks that we ultimately switched to hosting RTX-KG2c using a dedicated in-memory graph database engine, PloverDB (Sec. 2.2), that we developed. On the other hand, our standard procedure of hosting a Neo4j database server for RTX-KG2pre has been invaluable as a debugging aid and for developing graph queries and analysis workflows.

The performance of the RTX-KG2 build process has been parallelized where feasible. Although ETL-focused, open-source workflow tools such as Pentaho Kettle and Apache Hop are available, we selected Snakemake because of its seamless integration with Python and its parallelization support. Since RTX-KG2 was intended to be a large knowledge graph, the build process was designed to limit the amount of time it would take to build, especially by using the ETL process described above and by utilizing a parallel build system. With parallelization, the RTX-KG2pre build takes about 47 hours versus ∼74 hours without parallelization. The system has the capability to carry out a partial rebuild (skipping the most time-consuming updates of UMLS and SemMedDB) which can reduce rebuild time by ∼60%. In addition, Snakemake functionality along with the ETL-based approach allows for unchanged source databases to be ignored until the `Merge` step (see Fig. 10), which reduces build time. While building RTX-KG2 requires short-term access to a system with at least 256 GiB of physical memory, once RTX-KG2 is built, it can be hosted on a commodity server system (e.g., 64 GiB of memory; see Sec. A.3.1). New versions of RTX-KG2 are released at a rate of approximately once per month, with the median percentage increase in the number of edges per release being 0.8%, a relatively steady rate of growth consistent with the monthly rate of growth of abstracts in PubMed (0.6%).

**Figure 10:**
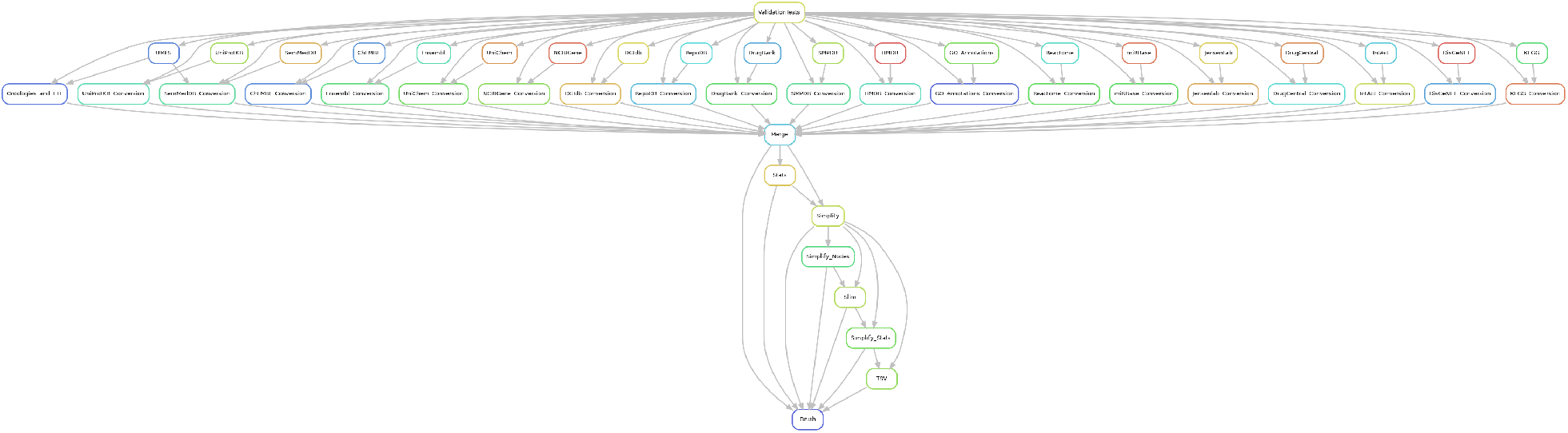
Flowchart of tasks for building RTX-KG2pre (the precursor stage of RTX-KG2) from 21 upstream knowledge-base distributions.

## 4 Conclusions

Despite the advances in the field outlined in Sec. 1, no open-source software toolkit was available that could integrate UMLS sources, SemMedDB, ChEMBL, DrugBank, SMPDB, Reactome, and 23 OBO Foundry ontologies (70 sources in all) into a single canonicalized knowledge graph based on the open-standard Biolink model as the semantic layer. To fill this gap and to provide a comprehensive knowledge-base to serve as as an efficient and accessible knowledge-substrate for a biomedical reasoning engine, we constructed RTX-KG2, comprising a set of ETL modules, an integration module, a REST API, and a parallel-capable build system that produces and hosts both pre-canonicalized (RTX-KG2pre) and canonicalized (RTX-KG2c) knowledge graphs for download and for querying. Quantitative usage information shows that RTX-KG2 is currently extensively used by multiple reasoning agents in the NCATS Biomedical Data Translator project, validating the ETL-focused, monolithic-graph, standards-based design philosophy that guided RTX-KG2’s development.

## 5 List of Abbreviations

ARAGORN: Autonomous Relay Agent for Generation Of Ranked Networks
ARAX: Autonomous Relay Agent X
ARS: Autonomous Relay System
AWS: Amazon Web Services
BTE: BioThings Explorer
D2J: direct-to-JSON method
EC2: Elastic Compute Cloud
ETL: extract–transform–load paradigm
GO: Gene Ontology
ICD: International Classification of Diseases
JSON: JavaScript Object Notation
KEGG: Kyoto Encyclopedia of Genes and Genomes
NCATS: National Center for Advancing Translational Sciences
NCBI: National Center for Biotechnology Information
OBO: Open Biomedical Ontologies
OWL: Web Ontology Language
RBM: RDF-based method
RDF: Resource Description Framework
REST: REpresentational State Transfer
RTX-KG2: Reasoning Tool X, Knowledge Graph Generation Two
RTX-KG2c: Reasoning Tool X, Knowledge Graph Generation Two, Canonicalized
RTX-KG2pre: Reasoning Tool X, Knowledge Graph Generation Two, Pre-canonicalization
S3: Simple Storage Service
SemMedDB: Semantic Medline Database
SMPDB: Small Molecule Pathway Database
SQL: Structured Query Language
Translator: NCATS Biomedical Data Translator
TRAPI: Translator Reasoner API
TSV: tab-separated value
TTL: Terse RDF Triple Language
UMLS: Unified Medical Language System
XML: eXtensible Markup Language
YAML: Yet Another Markup Language

(See also Table 2.1, Table 7, and Table S1).

## 6 Declarations

### 6.1 Ethics approval and consent to participate

Not applicable

### 6.2 Consent for publication

Not applicable

### 6.3 Availability of data and materials

The code supporting the conclusions of this article is available in the GitHub repository [github:RTXteam/RTX-KG2](https://github.com/RTXteam/RTX-KG2). Downloadable versions of RTX-KG2pre and RTX-KG2c are publicly available at [github:ncats/translator-lfs-artifacts](https://github.com/ncats/translator-lfs-artifacts/). The RTX-KG2 API is registered via the SmartAPI framework and can be reached at arax.rtx.ai/api/rtxkg2/v1.2/openapi.json.

The web browser user interface for querying RTX-KG2 is publicly available at arax.rtx.ai/kg2.

### 6.6 Authors’ contributions

Wrote the paper: ECW AKG LGK SAR MS ASH; designed the studies: SAR ECW AKG EWD DK; carried out computational work: ECW AKG SAR DK EWD FW LGK LA LM MS CM TSY JCR MS AT YC VF; evaluation, testing, and feedback: AKG ECW DK EWD JCR ASH SAR FW LGK CM LM MS LA TSY AT YC VF. All authors read and approved the final manuscript.

## 6.7 Acknowledgements

We thank Yao Yao, Zheng Liu, and Deqing Qu for technical assistance. We thank Chris Mungall, Tom Conlin, Matt Brush, Chunlei Wu, Harold Solbrig, Will Byrd, Michael Patton, Jim Balhoff, Chris Bizon, Deepak Unni, Richard Bruskiewich, Andrew Su, Kevin Xin, Mark Williams, and Jeff Henrikson for technical advice and/or feedback. We thank David Wishart and Carin Li for providing a download link for the SMPDB PubMed annotations and NCATS for help with hosting RTX-KG2 on GitHub. AKG gratefully acknowledges support from the ARCS Foundation. TSY gratefully acknowledges support from the Semiconductor Research Corporation.

## A Technical details of database construction

### A.1 Building RTX-KG2pre from upstream sources

The process by which the RTX-KG2 system builds its knowledge graph from its 70 sources—the first stage of which is diagrammed in Fig. 10)—begins by executing validation Python scripts (the “`validationTests`” task in Fig 10) that ensure that the identifiers used in the RTX-KG2 semantic layer are syntactically and semantically correct within the Biolink model. Next, the build process executes in parallel the 21 direct-to-JSON and RDF-based ETL scripts (see second and third rows in Fig. 10 and Sec. 2.1) to produce a total of 21 JSON files (20 via the direct-to-JSON method and one via the RDF-based method) in the RTX-KG2 knowledge graph schema (described in Sec. 2.3). Next, those JSON files are loaded and their object models are merged (via the “`Merge`” task) into a single self-contained graph that is then saved in an RTX-KG2-schema JSON file in which relationships consist of triples (subject, relation, object) where the relation is from any number of source-specific relationship vocabularies (totalling 1,228 source-specific relationship types in all). The RTX-KG2 object model is then simplified (via the “`Simplify`” task) by consolidating redundant relationships; a redundant relationship is where two or more sources assert the same triple (`ibuprofen`, `treats`, `headache`), in which case, the multiple relationships with identical triples are merged into a single relationship, with the multiple sources noted in the list-type `provided-by` relationship attribute (see Sec. 2.3). Also in the `Simplify` task, relationship types are simplified by mapping each of them to one of 77 standardized relationship types (called “predicates”) in the Biolink model. This mapping process is controlled by a locally maintained, rule-based system encoded in Yet Another Markup Language (YAML) that is cross-checked by a validation script against the Biolink model at build time. In that ruleset, some semantic loss of precision is allowed, such as mapping `DGIdb:partial_antagonist` to `biolink:decreases_activity of`, as a practical choice since the number of Biolink predicates (77) is much smaller than the number of total number of source relationship types. The `Simplify` task also maps source identifiers to Biolink “Information Resource” identifiers. In general, the `Simplify` task standardizes the graph with the Biolink model standards [49, 50]. We call the resulting graph *RTX-KG2pre* in order to emphasize that it is the precursor graph to the canonicalized RTX-KG2 graph (described below). In the final step of the build process, the nodes and edges of RTX-KG2pre are imported into a Neo4j graph database (for details, see Sec. A.3.2). During the RTX-KG2pre build, nodes (concepts) are assigned Biolink categories based on a rule-based system encoded in Yet Another Markup Language (YAML); at the start of each build, the YAML file is cross-checked by a validation script against known concept-to-category mappings in the Biolink model, to ensure correctness. The rule-based system allows for recursive assignment of a category to a class and all its subclasses, as well as to all concepts that have a specific CURIE prefix.

#### A.1.1 ARAX Node Synonymizer

As described in Sec. 2.2, in order to cluster concept identifiers into synonym groups (i.e., “canonicalize” the graph), the RTX-KG2 build system uses the ARAX [65] *Node Synonymizer* service, which takes into account four sources of evidence in the following order: (i) concept equivalence information obtained dynamically by querying a Translator web service API called the Standards and Reference Implementations (SRI) Node Normalizer ([github:TranslatorSRI/NodeNormalization](https://github.com/TranslatorSRI/NodeNormalization)); (ii) `biolink:same as` edges in RTX-KG2pre between RTX-KG2pre nodes; (iii) human-recognizable node (concept) name equivalence; and (iv) node semantic type compatibility. For dividing nodes into disjoint sets of equivalent nodes (i.e., Step 2 in Sec. 2.2), the Node Synonymizer goes through three passes of merging concepts in order to ensure that the partitioning of nodes is independent of the order in which the nodes are loaded into the Node Synonymizer. For choosing a canonical node for each concept cluster (i.e., Step 3 in Sec. 2.2), the Node Synonymizer uses a score-based system that flexibly enables incorporation of new heuristics. For the relatively small local table store for the ARAX Node Synonymizer, SQLite was chosen due to its combination of simplicity of deployment due to being serverless, its simplicity of use, and its faster response times vs. a network-based relational database management system solution. Performance of the Node Synonymizer is primarily tracked during the build process via a `problems.tsv` file, in which concept clusters that contain clashing categories (e.g., Drug and Disease) are recorded. The number of such “problem” clusters is monitored and any notable increases are manually debugged and corrected. While helpful, this performance metric relies on hard-coded definitions of which categories are considered clashing, does not catch potential merge errors that involve nodes of the same or compatible categories, and does not provide any insight into merge misses. In the future we plan to develop methods for more accurate performance assessment that do not have these limitations.

### A.2 Detailed schema for RTX-KG2

RTX-KG2pre and RTX-KG2c are serialized as JSON with the former’s schema being largely a superset of the latter’s. The JSON-serialized RTX-KG2pre has top-level keys `build`, `nodes`, and `edges`, with the `nodes` object containing a list of serialized objects for the concept nodes in the graph, and with `edges` containing a list of serialized objects for the subject-object-relationship triples in the graph. Under the `build` key, a JSON object stores information about the version of RTX-KG2pre and timestamp of the build. Under the `nodes` key is a list containing a JSON object for each node. Each node object contains 16 keys corresponding to the node property types in RTX-KG2pre, detailed in Table 4. The `id` node property contains a compact representation of the canonical uniform resource identifier, i.e., a CURIE [118]. The `category` property of a node describes the node’s semantic type, such as `biolink:Gene`. Similarly, the `edges` key is a list containing a JSON object for each edge, with the edge JSON object containing the keys indicated in Table 5. The `predicate` edge property contains the relationship type from the Biolink relationship hierarchy (which is called a “predicate” in the Biolink model), and the `relation` edge property contains the source relationship, such as `semmeddb:treats`.

**Table 4:** Node properties in RTX-KG2pre and RTX-KG2c

|  | KG2pre | KG2c |
| --- | --- | --- |
| all_categories |  | ✓ |
| all_names |  | ✓ |
| category | ✓ | ✓ |
| category_label | ✓ |  |
| creation_date | ✓ |  |
| deprecated | ✓ |  |
| description | ✓ | ✓ |
| equivalent_curies |  | ✓ |
| full_name | ✓ |  |
| has_biological_sequence | ✓ |  |
| id | ✓ | ✓ |
| iri | ✓ | ✓ |
| knowledge_source | ✓ |  |
| name | ✓ | ✓ |
| provided_by | ✓ |  |
| publications | ✓ | ✓ |
| replaced_by | ✓ |  |
| synonym | ✓ |  |
| update_date | ✓ |  |

**Table 5:** Edge properties in RTX-KG2pre and RTX-KG2c

|  | KG2pre | KG2c |
| --- | --- | --- |
| id | ✓ | ✓ |
| kg2_ids |  | ✓ |
| knowledge_source | ✓ | ✓ |
| negated | ✓ |  |
| object | ✓ | ✓ |
| predicate | ✓ | ✓ |
| predicate_label | ✓ |  |
| provided_by | ✓ |  |
| publications | ✓ | ✓ |
| publications_info | ✓ | ✓ |
| relation | ✓ |  |
| relation_label | ✓ |  |
| subject | ✓ | ✓ |
| update_date | ✓ |  |

The schema of the JSON serialization of RTX-KG2c is very close to that of RTX-KG2pre except that the former does not contain the top-level `build` key/object and, for each node object, RTX-KG2c contains some additional keys such as `equivalent_curies`, which enumerates the CURIEs of the nodes representing concepts that were semantically identified in the canonicalization step; `all_names`, which contains the `name` properties of the KG2pre nodes that were canonicalized together for the given KG2c node; and `all_categories`, which contains the `category` properties of the nodes that were canonicalized together for the given KG2c node. In addition, the `id`s of the corresponding KG2pre edges that each KG2c edge was created from are documented in the `kg2_ids` property.

### A.3 RTX-KG2 build system and software

In this section we describe the Snakemake-based build system for RTX-KG2 and the build system infrastructure requirements for building the RTX-KG2pre and RTX-KG2c graphs.

#### A.3.1 Requirements

The software for building RTX-KG2pre is designed to run in the Ubuntu Linux version 18.04 operating system on a dedicated system with at least 256 GiB of memory, 1 TiB of disk space in the root file system, ≥ 1 Gb/s networking, and at least 20 cores (we use an Amazon Web Services (AWS) Elastic Compute Cloud (EC2) instance of type `r5a.8xlarge`). The software for building RTX-KG2 makes use of AWS Simple Storage Service (S3) for network storage of both build artifacts and input knowledge source distribution files that cannot be retrieved by a scripted HTTP GET from their respective providers (see Table 6). These build files must be pre-staged in an AWS S3 bucket before the build process for RTX-KG2pre is started. For hosting RTX-KG2 in a Neo4j server, the system requirements are 64 GiB of system memory, 8 virtual CPUs, and ∼200 GiB of root filesystem storage (we use a `r5a.2xlarge` instance).

**Table 6:** Upstream source files that must be staged in S3 in order to build RTX-KG2

|  |  |  |
| --- | --- | --- |
| DrugBank | XML Download | Requires browser to download |
| RepoDB | TSV Download | Requires browser to download |
| SemMedDB | MySQL Download | Requires browser to download |
| SMPDB PubMed IDs | CSV Download | Obtained courtesy of Wishart Lab |
| UMLS Metathesaurus | ZIP Download | Requires browser to download |

**Table 7:**
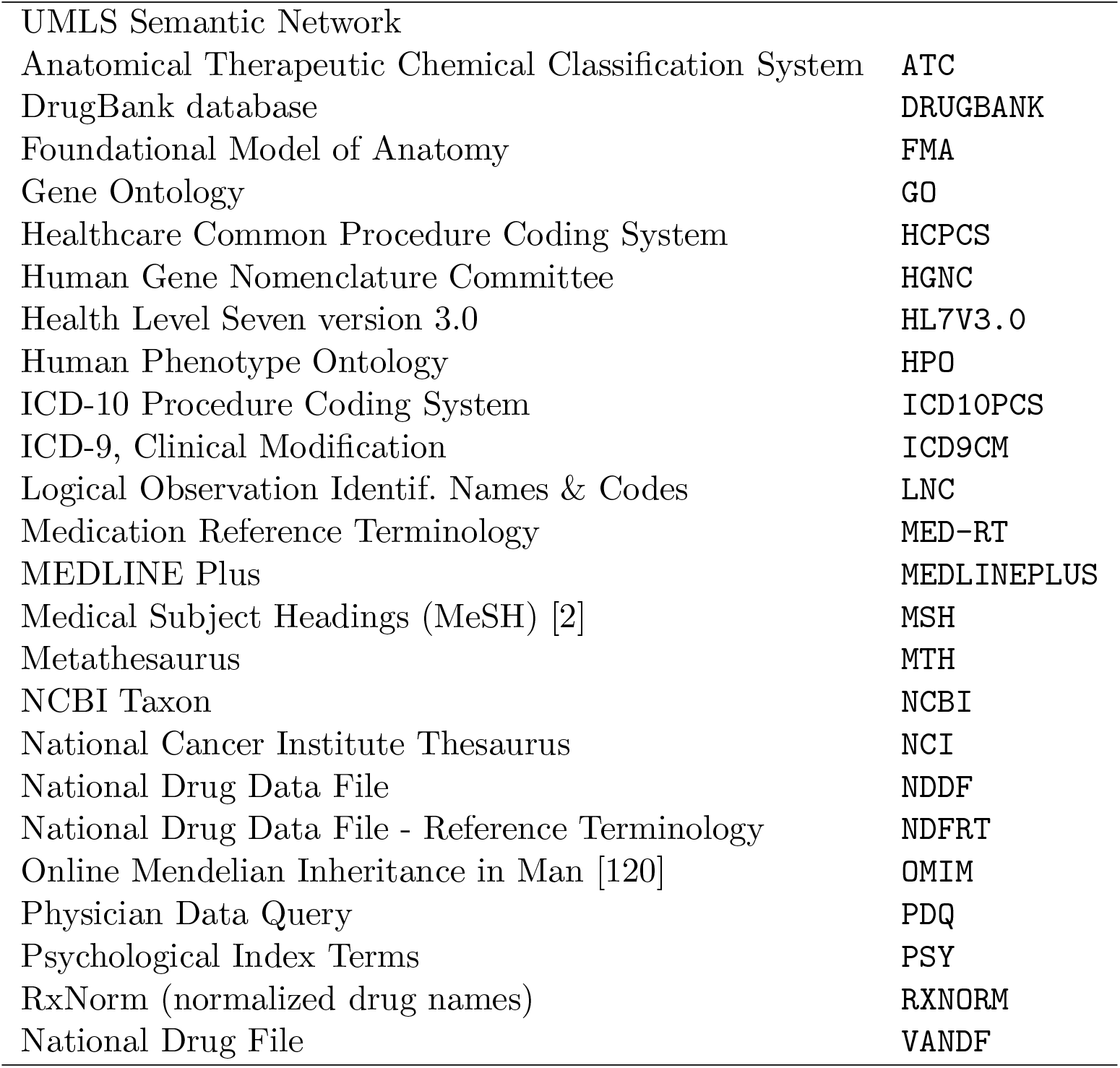
UMLS sources that are integrated into RTX-KG2. See Sec. 5 for definitions of abbreviations. See Sec. 3.4 regarding UMLS sources that could not be included due to licensing.

#### A.3.2 RTX-KG2 uses Snakemake for building RTX-KG2pre

RTX-KG2pre is built by a series of Python modules and bash scripts that extract, transform and load (ETL) 45 data downloads (corresponding to the rows of Table 1, with the “OBO Foundry” row counting for 21 separate downloads) from 24 source websites (Sec. 2.1) into a standardized property knowledge graph format integrated with the Biolink model as the semantic layer. To maximize the reproducibility of RTX-KG2 builds, the build system is fully automated, including scripts for (i) setting up and configuring the build system to run, (ii) downloading and transforming data, and (iii) exporting the final graph to the graph database Neo4j. RTX-KG2 utilizes the Snakemake [63] workflow management tool to schedule multicore execution of the RTX-KG2pre build process. In addition to reducing the computational costs of the build and the amount of time it takes to run, Snakemake increases modularity by enabling individual components (and their upstream dependencies) to be executed, when necessary. This is particularly useful for allowing failed builds to resume at the point of failure (via a so-called “partial” build), once the root cause (which could be a parsing error from an upstream ontology, for example) has been fixed.

The build process starts with parallel source extractions, in which all of the source databases are downloaded and prepared for the format that their respective conversion script uses. Then, each upstream source’s dataset is processed by a Python conversion module. This converts each source’s data into the RTX-KG2pre JSON format (Sec. 2.3). Once all of the upstream data sources are converted into their RTX-KG2pre JSON file, a module merges all of them into a cohesive graph, such that no two nodes have the same CURIE ID. One of the challenges in this step is when different upstream sources provide different names for the same concept CURIE; the RTX-KG2pre build system addresses such name conflicts by having a defined order of precedence of upstream sources. After the merge step, edge source relation types (as described in Sec. A.1) are each mapped to one of 77 predicate types in the Biolink predicate hierarchy (see Sec. A.1), and redundant edges (same combination of subject node ID, object node ID, and Biolink predicate) are coalesced, with source relation information and source provenance information added to lists in the coalesced edge. The graph is then serialized as JSON (see Sec. 2.3) and to TSV format. The build artifacts, including the unprocessed and processed JSON files and the TSV files, are uploaded into an AWS S3 bucket. RTX-KG2pre is then hosted in Neo4j on a smaller AWS instance (see Sec. A.3.1); the Neo4j endpoint is mainly used for debugging.

#### A.3.3 umls2rdf and owltools

In the RTX-KG2pre build process, the 26 UMLS sources are ingested as TTL files that are generated in the extraction stage of the build process from the Rich Release Format (RRF [68]) UMLS distribution using two software programs, Metamorphosys [119] (to load the RRF files into the relational database system, MySQL) and umls2rdf [29] (to extract TTL files Sec. A.3.3). Thus, a local MySQL database is used as an intermediate data source in the build process, from which TTL files are generated via `umls2rdf`. The build system uses the `owltools` software program ([github:owlcollab/owltools](https://github.com/owlcollab/owltools)) to convert biomedical ontologies (see Table 1 and Table S1) in OWL format and the UMLS TTL files into OBO (Open Biological and Biomedical Ontology) JSON format for processing. The ontologies in OBO-JSON format are then loaded using the Python package `ontobio` (which, in turn, is based on the NetworkX graph library [121]) and processed/merged together, enabling use of cross-ontology axioms in determining concept semantic types.

## B Supplementary Material

**Table S1:** Ontologies from the OBO Foundry that are included in RTX-KG2. Basic Formal Ontology BFO

|  |  |
| --- | --- |
| Basic Formal Ontology | BFO |
| Chemical Entities of Biological Interest (ChEBI) | CHEBI [122] |
| Gene Ontology, with external relationships | go-plus |
| Relation Ontology | RO |
| Uberon multi-species anatomy ontology, extended with external relationships | UBERON |
| Foundational Model of Anatomy | FMA |
| Dictyostelium discoideum anatomy | DDANAT [123–127] |
| Cell Ontology | CL [128] |
| Food Ontology | FOODON |
| Human Developmental Anatomy, abstract | EHDAA2 [129] |
| Biological Spatial Ontology | BSPO |
| Human Phenotype Ontology | HPO |
| Neuro Behavior Ontology | NBO |
| NCBI organismal classification, taxslim subset | ncbitaxon [103] |
| Phenotype and Trait Ontology | PATO |
| Mondo Disease Ontology | MONDO |
| Disease Ontology | DO |
| Protein Ontology | PRO [130] |
| Interaction Network Ontology | INO |
| Genomic Epidemiology Ontology | GENEPIO |
| Molecular Interactions Controlled Vocabulary | MI |

## Footnotes

1 An example identifier type prioritization would be for the semantic type “gene”, to prefer (from most to least preferred) identifier types from Ensembl Gene, NCBI Gene, and Human Gene Nomenclature Committee (HGNC).

2 This will be transitioning to the original predicate property in the next release of RTX-KG2, for compatibility with recent changes in the Biolink standard.

3 [github:NCATSTranslator/testing/ars-requests/not-none/1.2](https://github.com/NCATSTranslator/testing/tree/main/ars-requests/not-none/1.2), accessed on June 23, 2022

4 Note however, that one API is used in constructing RTX-KG2; see Sec. 2.1.

## References

[1] Philip R O Payne. Chapter 1: Biomedical Knowledge Integration. PLOS Comput Biol, 8(12):e1002826, 2012.

[2] F B Rogers. Medical Subject Headings. Bulletin of the Medical Library Association, 51(1):114–116, 1963.

[3] A W Forrey, C J McDonald, G DeMoor, et al. Logical observation identifier names and codes (LOINC) database: a public use set of codes and names for electronic reporting of clinical laboratory test results. Clin Chem, 42(1):81–90, 1996.

[4] Y A Lussier, D J Rothwell, and R A Côté. The SNOMED model: a knowledge source for the controlled terminology of the computerized patient record. Methods Inf Med, 37(2):161–164, 1998.

[5] E G Brown, L Wood, and S Wood. The medical dictionary for regulatory activities (MedDRA). Drug Safety, 20(2):109–117, 1999.

[6] Stuart J Nelson, Kelly Zeng, John Kilbourne, et al. Normalized names for clinical drugs: RxNorm at 6 years. J Am Med Inform Assoc, 18(4):441–448, 2011.

[7] B L Humphreys, D A Lindberg, H M Schoolman, and G O Barnett. The Unified Medical Language System: an informatics research collaboration. J Am Med Inform Assoc, 5(1):1–11, 1998.

[8] Jonathan Bard, Seung Y Rhee, and Michael Ashburner. An ontology for cell types. Genome Biol, 6(2):R21, 2005.

[9] D Brickley and R V Guha. Resource description framework (RDF) schema specification. Technical Report 19990303, World Wide Web Consortium, Cambridge, MA, USA, 1999. URL: https://www.w3.org/TR/1999/PR-rdf-schema-19990303/.

[10] Sean Bechhofer, Frank van Harmelen, Jim Hendler, et al. Owl web ontology language reference. Technical Report 20040210, World Wide Web Consortium, Cambridge, MA, USA, 2004. URL: https://www.w3.org/TR/2004/REC-owl-ref-20040210/.

[11] M. Kanehisa. KEGG: Kyoto encyclopedia of genes and genomes. Nucleic Acids Res, 28(1):27–30, 2000. doi:10.1093/nar/28.1.27.

[12] Sunghwan Kim, Jie Chen, Tiejun Cheng, et al. PubChem in 2021: new data content and improved web interfaces. Nucleic Acids Res, 49(D1):D1388–D1395, 2021.

[13] D. S. Wishart. DrugBank: a comprehensive resource for in silico drug discovery and exploration. Nucleic Acids Res, 34(90001):D668–D672, 2006. doi:10.1093/nar/gkj067.

[14] David Mendez, Anna Gaulton, A Patrícia Bento, et al. ChEMBL: towards direct deposition of bioassay data. Nucleic Acids Res, 47(D1):D930–D940, 2018. doi:10.1093/nar/gky1075.

[15] Alex Bateman, Maria-Jesus Martin, Sandra Orchard, et al. UniProt: the Universal Protein Knowledgebase in 2021. Nucleic Acids Res, 49(D1):D480–D489, 2020. doi:10.1093/nar/gkaa1100.

[16] Alex Frolkis, Craig Knox, Emilia Lim, et al. SMPDB: The Small Molecule Pathway Database. Nucleic Acids Res, 38(Suppl 1):D480–D487, 2009. doi:10.1093/nar/gkp1002.

[17] Timothy Jewison, Yilu Su, Fatemeh Miri Disfany, et al. SMPDB 2.0: Big improvements to the small molecule pathway database. Nucleic Acids Res, 42(D1):D478–D484, 2013. doi:10.1093/nar/gkt1067.

[18] Antonio Fabregat, Florian Korninger, Guilherme Viteri, et al. Reactome graph database: Efficient access to complex pathway data. PLOS Comput Biol, 14(1):e1005968, 2018. doi:10.1371/journal.pcbi.1005968.

[19] Thomas C Rindflesch and Marcelo Fiszman. The interaction of domain knowledge and linguistic structure in natural language processing: interpreting hypernymic propositions in biomedical text. J Biomed Inform, 36(6):462–477, 2003.

[20] Sergey Goryachev, Margarita Sordo, and Qing T Zeng. A suite of natural language processing tools developed for the I2B2 project. American Medical Informatics Association Symposium proceedings, 2006:931–931, 2006.

[21] Marco A Valenzuela-Escárcega, Özgün Babur, Gus Hahn-Powell, et al. Large-scale automated machine reading discovers new cancer-driving mechanisms. Database, 2018.

[22] Rebecca Sharp, Adarsh Pyarelal, Benjamin Gyori, et al. Eidos, INDRA, & Delphi: from free text to executable causal models. In Proceedings of the 2019 Conference of the North American Chapter of the Association for Computational Linguistics (Demonstrations), 2019.

[23] Rui Xing, Jie Luo, and Tengwei Song. BioRel: towards large-scale biomedical relation extraction. BMC Bioinformatics, 21(16):543, 2020.

[24] Mila Glavaški and Lazar Velicki. Humans and machines in biomedical knowledge curation: hypertrophic cardiomyopathy molecular mechanisms’ representation. BioData Min, 14(1):45, 2021.

[25] National Library of Medicine (US). Pubmed [internet], 1964. URL: https://www.ncbi.nlm.nih.gov/pubmed/.

[26] H. Kilicoglu, D. Shin, M. Fiszman, et al. SemMedDB: a PubMed-scale repository of biomedical semantic predications. Bioinformatics, 28(23):3158–3160, 2012. doi:10.1093/bioinformatics/bts591.

[27] Barry Smith, Werner Ceusters, Bert Klagges, et al. Relations in biomedical ontologies. Genome Biol, 6(5):R46, 2005.

[28] Elena Beisswanger, Stefan Schulz, Holger Stenzhorn, and Udo Hahn. BioTop: an upper domain ontology for the life sciences. Appl Ontol, 3(4):205–212, 2008.

[29] Mark A Musen, Natalya F Noy, Nigam H Shah, et al. The National Center for Biomedical Ontology. J Am Med Inform Assoc, 19(2):190–195, 2012.

[30] Michel Dumontier, Christopher JO Baker, Joachim Baran, et al. The Semanticscience Integrated Ontology (SIO) for biomedical research and knowledge discovery. J Biomed Semantics, 5(1):14, 2014.

[31] Rebecca Jackson, Nicolas Matentzoglu, James A Overton, et al. OBO Foundry in 2021: operationalizing open data principles to evaluate ontologies. Database, 2021. doi:10.1093/database/baab069.

[32] Tunca Doğan, Heval Atas, Vishal Joshi, et al. CROssBAR: comprehensive resource of biomedical relations with deep learning applications and knowledge graph representations. bioRxiv, 2020. doi:10.1101/2020.09.14.296889.

[33] Pablo Pareja-Tobes, Raquel Tobes, Marina Manrique, et al. Bio4j: a high-performance cloud-enabled graph-based data platform. bioRxiv, 2015. doi:10.1101/016758.

[34] Aaron Birkland and Golan Yona. BIOZON: a system for unification, management and analysis of heterogeneous biological data. BMC Bioinformatics, 7(1):70, 2006.

[35] Antonino Fiannaca, Massimo La Rosa, Laura La Paglia, et al. Biographdb: a new graphdb collecting heterogeneous data for bioinformatics analysis. In Eighth International Conference on Bioinformatics, Biocomputational Systems and Biotechnologies, Wilmington, 2016. IARIA.

[36] Daniel Scott Himmelstein, Antoine Lizee, Christine Hessler, et al. Systematic integration of biomedical knowledge prioritizes drugs for repurposing. eLife, 6:e26726, 2017. doi:10.7554/eLife.26726.

[37] Sergio Baranzini, Sui Huang, Sharat Israni, et al. Scalable precision medicine knowledge engine, 2021. Accessed: 2021-06-01. URL: https://spoke.ucsf.edu.

[38] Geoffrey Sanders, Roger Pearce, and Sergio E. Baranzini. Topological analysis of the SPOKE graph. Technical report, U. S. Department of Energy, 2020. doi:10.2172/1669224.

[39] Yi Liu, Benjamin Elsworth, Pau Erola, et al. EpiGraphDB: a database and data mining platform for health data science. Bioinformatics, 2020.

[40] Vassilis N Ioannidis, Da Zheng, and George Karypis. Few-shot link prediction via graph neural networks for covid-19 drug-repurposing. arXiv preprint arXiv:2007.10261, 2020.

[41] Michel Dumontier, Alison Callahan, Jose Cruz-Toledo, et al. Bio2RDF release 3: a larger connected network of linked data for the life sciences. In Proceedings of the 2014 International Conference on Posters & Demonstrations Track, volume 1272, pages 401–404. Citeseer, 2014.

[42] Kevin M Livingston, Michael Bada, William A Baumgartner, and Lawrence E Hunter. KaBOB: ontology-based semantic integration of biomedical databases. BMC Bioinformatics, 16(1):126, 2015.

[43] Yong Zhang, Ming Sheng, Rui Zhou, et al. HKGB: an inclusive, extensible, intelligent, semi-auto-constructed knowledge graph framework for healthcare with clinicians’ expertise incorporated. Inf Process Manag, 57(6):102324, 2020. doi:https://doi.org/10.1016/j.ipm.2020.102324.

[44] Kenneth Morton, Patrick Wang, Chris Bizon, et al. ROBOKOP: an abstraction layer and user interface for knowledge graphs to support question answering. Bioinformatics, 35(24):5382–5384, 2019.

[45] Karamarie Fecho, Chris Bizon, Frederick Miller, et al. A biomedical knowledge graph system to propose mechanistic hypotheses for real-world environmental health observations: cohort study and informatics application. JMIR Med Inform, 9(7):e26714, 2021. doi:10.2196/26714.

[46] Jiwen Xin, Cyrus Afrasiabi, Sebastien Lelong, et al. Cross-linking BioThings APIs through JSON-LD to facilitate knowledge exploration. BMC Bioinformatics, 19(1):30, 2018.

[47] William E. Byrd, Gregory Rosenblatt, Michael John Patton, et al. mediKanren: a system for bio-medical reasoning. In Proceedings of the 2020 ACM SIGPLAN International Conference on Functional Programming, 2020.

[48] Chris Mungall, Hirokazu Chiba, Shuichi Kawashima, et al. Logic programming for the biomedical sciences, 2020. doi:10.37044/osf.io/km9ux.

[49] Justin Reese, Deepak Unni, Tiffany J Callahan, et al. KG-COVID-19: a framework to produce customized knowledge graphs for COVID-19 response. bioRxiv, 2020.

[50] Richard Bruskiewich, Deepak Unni, Chris Mungall, et al. biolink/biolink-model: 2.0.0, 2021. doi:10.5281/ZENODO.4895425.

[51] Deepak R Unni, Sierra AT Moxon, Michael Bada, et al. Biolink model: a universal schema for knowledge graphs in clinical, biomedical, and translational science. Clin Transl Sci, 2022.

[52] Biomedical Data Translator Consortium. Toward a universal biomedical data translator. Clin Transl Sci, 12(2):86–90, 2019.

[53] Julie A McMurry, Sebastian Köhler, Nicole L Washington, et al. Navigating the phenotype frontier: The Monarch Initiative. Genetics, 203(4):1491–1495, 2016. doi:10.1534/genetics.116.188870.

[54] Christopher J Mungall, Julie A McMurry, Sebastian Köhler, et al. The Monarch Initiative: an integrative data and analytic platform connecting phenotypes to genotypes across species. Nucleic Acids Res, 45(D1):D712–D722, 2017.

[55] Kent A Shefchek, Nomi L Harris, Michael Gargano, et al. The Monarch Initiative in 2019: an integrative data and analytic platform connecting phenotypes to genotypes across species. Nucleic Acids Res, 48(D1):D704–D715, 2019. doi:10.1093/nar/gkz997.

[56] Luis Galárraga, Geremy Heitz, Kevin Murphy, and Fabian M Suchanek. Canonicalizing open knowledge bases. In Proceedings of the 23rd ACM International Conference on Conference on Information and Knowledge Management, pages 1679–1688, 2014.

[57] Antonio Messina, Haikal Pribadi, Jo Stichbury, et al. BioGrakn: a knowledge graph-based semantic database for biomedical sciences. In Leonard Barolli and Olivier Terzo, editors, Complex, Intelligent, and Software Intensive Systems, pages 299–309. Springer International Publishing, 2018.

[58] Andra Waagmeester, Gregory Stupp, Sebastian Burgstaller-Muehlbacher, et al. Science forum: Wikidata as a knowledge graph for the life sciences. eLife, 9:e52614, 2020. doi:10.7554/eLife.52614.

[59] Stephen Ramsey, David Koslicki, Yao Yao, et al. RTXteam/RTX: Initial proof-of-concept software version from November 2017, 2018. doi:10.5281/ZENODO.1185486.

[60] Christopher J. Mungall, Julie A. McMurry, Sebastian Köhler, et al. The Monarch Initiative: an integrative data and analytic platform connecting phenotypes to genotypes across species. Nucleic Acids Res, 45(D1):D712–D722, 2016. doi:10.1093/nar/gkw1128.

[61] Ben Elsworth. EpigraphDB, 2021. doi:10.5281/ZENODO.4534128.

[62] Tiffany J. Callahan, Ignacio J. Tripodi, Lawrence E. Hunter, and William A. Baumgartner. A framework for automated construction of heterogeneous large-scale biomedical knowledge graphs. bioRxiv, 2020. doi:10.1101/2020.04.30.071407.

[63] Johannes Köster and Sven Rahmann. Snakemake—a scalable bioinformatics workflow engine. Bioinformatics, 28(19):2520–2522, 2012.

[64] Amrapali Zaveri, Shima Dastgheib, Chunlei Wu, et al. smartAPI: towards a more intelligent network of web APIs. In Eva Blomqvist, Diana Maynard, Aldo Gangemi, et al., editors, The Semantic Web, pages 154–169. Springer International Publishing, 2017.

[65] Amy K. Glen, Chunyu Ma, Luis Mendoza, et al. ARAX: a graph-based modular reasoning tool for translational biomedicine. bioRxiv, 2022. doi:10.1101/2022.08.12.503810.

[66] Richard D Hipp. SQLite, 2020. URL: https://www.sqlite.org/index.html.

[67] Fabien Gandon, Guus Schreiber, and Dave Beckett. RDF 1.1 XML Syntax. Technical Report 20140225, World Wide Web Consortium, Cambridge, MA, USA, 2014. URL: http://www.w3.org/TR/2014/REC-rdf-syntax-grammar-20140225/.

[68] UMLS Team. UMLS Reference Manual, chapter 3. National Library of Medicine (US), Bethesda, 2009. URL: https://www.ncbi.nlm.nih.gov/books/NBK9685.

[69] Mark Davies, Micha-l Nowotka, George Papadatos, et al. ChEMBL web services: streamlining access to drug discovery data and utilities. Nucleic Acids Res, 43(W1):W612–W620, 2015. doi:10.1093/nar/gkv352.

[70] Sharon L Freshour, Susanna Kiwala, Kelsy C Cotto, et al. Integration of the Drug–Gene Interaction Database (DGIdb 4.0) with open crowdsource efforts. Nucleic Acids Res, 49(D1):D1144–D1151, 2020. doi:10.1093/nar/gkaa1084.

[71] Janet Piñero, Juan Manuel Ramírez-Anguita, Josep Saüch-Pitarch, et al. The DisGeNET knowledge platform for disease genomics: 2019 update. Nucleic Acids Res, 2019. doi:10.1093/nar/gkz1021.

[72] Sorin Avram, Cristian G Bologa, Jayme Holmes, et al. DrugCentral 2021 supports drug discovery and repositioning. Nucleic Acids Res, 49(D1):D1160–D1169, 2020. doi:10.1093/nar/gkaa997.

[73] Andrew D Yates, Premanand Achuthan, Wasiu Akanni, et al. Ensembl 2020. Nucleic Acids Res, 2019. doi:10.1093/nar/gkz966.

[74] James Malone, Ele Holloway, Tomasz Adamusiak, et al. Modeling sample variables with an Experimental Factor Ontology. Bioinformatics, 26(8):1112–1118, 2010.

[75] Seth Carbon, Eric Douglass, Benjamin M Good, et al. The Gene Ontology resource: enriching a GOld mine. Nucleic Acids Res, 49(D1):D325–D334, 2020. doi:10.1093/nar/gkaa1113.

[76] Michael Ashburner, Catherine A. Ball, Judith A. Blake, et al. Gene Ontology: tool for the unification of biology. Nat Genet, 25(1):25–29, 2000. doi:10.1038/75556.

[77] D. S. Wishart, D. Tzur, C. Knox, et al. HMDB: the human metabolome database. Nucleic Acids Res, 35(Database):D521–D526, 2007. doi:10.1093/nar/gkl923.

[78] D. S. Wishart, C. Knox, A. C. Guo, et al. HMDB: a knowledgebase for the human metabolome. Nucleic Acids Res, 37(Database):D603–D610, 2009. doi:10.1093/nar/gkn810.

[79] David S. Wishart, Timothy Jewison, An Chi Guo, et al. HMDB 3.0—the human metabolome database in 2013. Nucleic Acids Res, 41(D1):D801–D807, 2012. doi:10.1093/nar/gks1065.

[80] David S Wishart, Yannick Djoumbou Feunang, Ana Marcu, et al. HMDB 4.0: the human metabolome database for 2018. Nucleic Acids Res, 46(D1):D608–D617, 2017. doi:10.1093/nar/gkx1089.

[81] H. Hermjakob. IntAct: an open source molecular interaction database. Nucleic Acids Res, 32(90001):452D–455, 2004. doi:10.1093/nar/gkh052.

[82] S. Kerrien, B. Aranda, L. Breuza, et al. The IntAct molecular interaction database in 2012. Nucleic Acids Res, 40(D1):D841–D846, 2011. doi:10.1093/nar/gkr1088.

[83] Sune Pletscher-Frankild, Albert Pallejà, Kalliopi Tsafou, et al. DISEASES: text mining and data integration of disease–gene associations. Methods, 74:83–89, 2015. doi:10.1016/j.ymeth.2014.11.020.

[84] Minoru Kanehisa. Toward understanding the origin and evolution of cellular organisms. Protein Sci, 28(11):1947–1951, 2019. doi:10.1002/pro.3715.

[85] Minoru Kanehisa, Miho Furumichi, Yoko Sato, et al. KEGG: integrating viruses and cellular organisms. Nucleic Acids Res, 49(D1):D545–D551, 2020. doi:10.1093/nar/gkaa970.

[86] S. Griffiths-Jones. The microRNA registry. Nucleic Acids Res, 32(90001):109D–111, 2004. doi:10.1093/nar/gkh023.

[87] S. Griffiths-Jones. miRBase: microRNA sequences, targets and gene nomenclature. Nucleic Acids Res, 34(90001):D140–D144, 2006. doi:10.1093/nar/gkj112.

[88] S. Griffiths-Jones, H. K. Saini, S. van Dongen, and A. J. Enright. miRBase: tools for microRNA genomics. Nucleic Acids Res, 36(Database):D154–D158, 2007. doi:10.1093/nar/gkm952.

[89] A. Kozomara and S. Griffiths-Jones. miRBase: integrating microRNA annotation and deep-sequencing data. Nucleic Acids Res, 39(Database):D152–D157, 2010. doi:10.1093/nar/gkq1027.

[90] Ana Kozomara, Maria Birgaoanu, and Sam Griffiths-Jones. miRBase: from microRNA sequences to function. Nucleic Acids Res, 47(D1):D155–D162, 2018. doi:10.1093/nar/gky1141.

[91] NCBI Resource Coordinators. Database resources of the National Center for Biotechnology Information. Nucleic Acids Res, 44(D1):D7–D19, 2015. doi:10.1093/nar/gkv1290.

[92] S. S. Weinreich, R. Magnon, J. J. Sikkens, et al. Orphanet: een Europese database over zeldzame ziekten [Orphanet: a European database for rare diseases]. Nederlands tijdschrift voor geneeskunde, 152(9):518–519, 2008. URL: https://pubmed.ncbi.nlm.nih.gov/18389888/.

[93] Allison Pon, Timothy Jewison, Yilu Su, et al. Pathways with PathWhiz. Nucleic Acids Res, 43(W1):W552–W559, 2015. doi:10.1093/nar/gkv399.

[94] Miguel Ramirez-Gaona, Ana Marcu, Allison Pon, et al. A web tool for generating high quality machine-readable biological pathways. J Vis Exp, 120, 2017. doi:10.3791/54869.

[95] David S Wishart, Carin Li, Ana Marcu, et al. PathBank: a comprehensive pathway database for model organisms. Nucleic Acids Res, 48(D1):D470–D478, 2019. doi:10.1093/nar/gkz861.

[96] Bijay Jassal, Lisa Matthews, Guilherme Viteri, et al. The Reactome pathway knowledgebase. Nucleic Acids Res, 2019. doi:10.1093/nar/gkz1031.

[97] O. Bodenreider. The unified medical language system (UMLS): integrating biomedical terminology. Nucleic Acids Res, 32(90001):267D–270, 2004. doi:10.1093/nar/gkh061.

[98] Jon Chambers, Mark Davies, Anna Gaulton, et al. UniChem: a unified chemical structure cross-referencing and identifier tracking system. J Cheminform, 5(1), 2013. doi:10.1186/1758-2946-5-3.

[99] World Wide Web Consortium et al. RDF 1.1 Turtle: terse RDF triple language. Technical Report 20140225, World Wide Web Consortium, Cambridge, MA, USA, 2014. URL: https://www.w3.org/TR/turtle/.

[100] Drashtti Vasant, Laetitia Chanas, James Malone, et al. Ordo: an ontology connecting rare disease, epidemiology and genetic data. In Proceedings of ISMB, volume 30, 2014.

[101] Fatima Zohra Smaili, Xin Gao, and Robert Hoehndorf. Formal axioms in biomedical ontologies improve analysis and interpretation of associated data. Bioinformatics, 36(7):2229–2236, 2019. doi:10.1093/bioinformatics/btz920.

[102] Barry Smith and Werner Ceusters. Ontological realism: A methodology for coordinated evolution of scientific ontologies. Appl Ontol, 5(3-4):139–188, 2010.

[103] Conrad L Schoch, Stacy Ciufo, Mikhail Domrachev, et al. NCBI Taxonomy: a comprehensive update on curation, resources and tools. Database, 2020. doi:10.1093/database/baaa062.

[104] Roy Thomas Fielding. REST: Architectural Styles and the Design of Network-based Software Architectures. Doctoral dissertation, University of California, Irvine, 2000. URL: http://www.ics.uci.edu/~fielding/pubs/dissertation/top.htm.

[105] Meghamala Sinha and Stephen A Ramsey. Using a general prior knowledge graph to improve data-driven causal network learning. In AAAI Spring Symposium: Combining Machine Learning with Knowledge Engineering, 2021.

[106] Yodsawalai Chodpathumwan, Arash Termehchy, Stephen A. Ramsey, et al. Structural generalizability: the case of similarity search. In Proceedings of the 2021 International Conference on Management of Data, SIGMOD/PODS ‘21, page 326–338, New York, NY, USA, 2021. Association for Computing Machinery. doi:10.1145/3448016.3457316.

[107] Finn Womack, Jason McClelland, and David Koslicki. Leveraging distributed biomedical knowledge sources to discover novel uses for known drugs. bioRxiv, 2019. doi:10.1101/765305.

[108] Deepak Unni and Kent Shefchek. SRI Reference KG, 2022. URL: https://github.com/Knowledge-Graph-Hub/sri-reference-kg.

[109] Melanie Courtot, Frank Gibson, Allyson Lister, et al. MIREOT: the Minimum Information to Reference an External Ontology Term. Nature Precedings, 2009.

[110] Leslie F Sikos and Dean Philp. Provenance-aware knowledge representation: a survey of data models and contextualized knowledge graphs. Data Sci Eng, 5(3):293–316, 2020.

[111] Deepak Unni, Richard Bruskiewich, Lance Hannestad, et al. Knowledge graph exchange library, 2021. URL: https://github.com/biolink/kgx.

[112] Mark Steyvers and Joshua B Tenenbaum. The large-scale structure of semantic networks: statistical analyses and a model of semantic growth. Cogn Sci, 29(1):41–78, 2005.

[113] Yuehang Ding, Hongtao Yu, Ruiyang Huang, and Yunjie Gu. Complex network based knowledge graph ontology structure analysis. In 2018 1st IEEE International Conference on Hot Information-Centric Networking (HotICN). IEEE, 2018. doi:10.1109/hoticn.2018.8606002.

[114] Jane Fedorowicz. A Zipfian model of an automatic bibliographic system: an application to MEDLINE. J Am Soc Inf Sci, 33(4):223–232, 1982. doi:10.1002/asi.4630330406.

[115] Leila Ranandeh Kalankesh, Robert Stevens, and Andy Brass. The language of gene ontology: a Zipf’s law analysis. BMC Bioinformatics, 13(1):127, 2012.

[116] Lawrence Page, Sergey Brin, Rajeev Motwani, and Terry Winograd. The PageRank citation ranking: bringing order to the web. Technical report, Stanford InfoLab, 1999.

[117] Nadime Francis, Alastair Green, Paolo Guagliardo, et al. Cypher: an evolving query language for property graphs. In Proceedings of the 2018 International Conference on Management of Data, pages 1433–1445, 2018.

[118] Mark Birbeck and Shane McCarron. CURIE syntax 1.0: a syntax for expressing compact URIs. Technical Report 20101216, World Wide Web Consortium, Cambridge, MA, USA, 2010. URL: https://www.w3.org/TR/2010/NOTE-curie-20101216/.

[119] Olivier Bodenreider. The Unified Medical Language System (UMLS): integrating biomedical terminology. Nucleic Acids Res, 32(Database issue):D267–70, 2004.

[120] Victor A McKusick. Mendelian Inheritance in Man and its online version, OMIM. Am J Hum Genet, 80(4):588–604, 2007.

[121] Aric A. Hagberg, Daniel A. Schult, and Pieter J. Swart. Exploring network structure, dynamics, and function using NetworkX. In Gäel Varoquaux, Travis Vaught, and Jarrod Millman, editors, Proceedings of the 7th Python in Science Conference, pages 11 – 15, Pasadena, CA USA, 2008.

[122] Janna Hastings, Gareth Owen, Adriano Dekker, et al. ChEBI in 2016: Improved services and an expanding collection of metabolites. Nucleic Acids Res, 44(D1):D1214–D1219, 2015. doi:10.1093/nar/gkv1031.

[123] Petra Fey, Robert J. Dodson, Siddhartha Basu, and Rex L. Chisholm. One stop shop for everything dictyostelium: dictyBase and the Dicty Stock Center in 2012. In Methods in Molecular Biology, pages 59–92. Humana Press, 2013. doi:10.1007/978-1-62703-302-2_4.

[124] Siddhartha Basu, Petra Fey, Yogesh Pandit, et al. dictyBase 2013: integrating multiple dictyostelid species. Nucleic Acids Res, 41(D1):D676–D683, 2012. doi:10.1093/nar/gks1064.

[125] Petra Fey, Pascale Gaudet, Tomaz Curk, et al. dictyBase—a dictyostelium bioinformatics resource update. Nucleic Acids Res, 37(Suppl 1):D515–D519, 2008. doi:10.1093/nar/gkn844.

[126] R. L. Chisholm. dictyBase, the model organism database for dictyostelium discoideum. Nucleic Acids Res, 34(90001):D423–D427, 2006. doi:10.1093/nar/gkj090.

[127] L. Kreppel. dictyBase: a new dictyostelium discoideum genome database. Nucleic Acids Res, 32(90001):332D–333, 2004. doi:10.1093/nar/gkh138.

[128] Chris Mungall, Shawn Tan, Nicole Vasilevsky, et al. obophenotype/cell-ontology: 2021-04-22 release, 2021. doi:10.5281/ZENODO.592969.

[129] Jonathan Bard. A new ontology (structured hierarchy) of human developmental anatomy for the first 7 weeks (Carnegie stages 1-20). J Anat, 221(5):406–416, 2012. doi:10.1111/j.1469-7580.2012.01566.x.

[130] Chuming Chen, Hongzhan Huang, Karen E. Ross, et al. Protein ontology on the semantic web for knowledge discovery. Sci Data, 7(1), 2020. doi:10.1038/s41597-020-00679-9.

